# The Effects of Early Life Pain and Juvenile Fear Conditioning on CRF-Receptor Expression in the Amygdala and Hypothalamus

**DOI:** 10.1101/2025.05.29.656935

**Authors:** Michael A. Burman, Jared T. Zuke

## Abstract

Early life pain and stress have lasting consequences on nervous system development that can interact with later stress or trauma to create a susceptibility to fear, anxiety, depression and chronic pain among other psychological disorders. Recent work has identified changes in corticotropin releasing factor signaling in limbic system structures, such as the amygdala and hypothalamus, as a key mechanism behind these changes – albeit in a sex-dependent manner. CRF has two major receptors, CRHR1 and CRHR2 which have also been shown to play key roles in fear and pain expression. The current work examines the effects of early life pain designed to mimic the neonatal medical trauma that occurs in the Neonatal Intensive Care Unit (NICU), paired with a juvenile trauma in the form of fear conditioning, on expression of *crhr1* and *crhr2* mRNA in the central (CeA) and basolateral (BLA) amygdala as well as the paraventricular (PVN) and ventromedial (VMH) hypothalamus. While prior work has demonstrated that early life pain significantly impacts expression of the CRF ligand, the current data demonstrate that early life pain and fear conditioning largely fail to affect CRH receptor expression in the amygdala. Modest changes in expression to *crhr2* in a sex and region-specific manner were observed in the hypothalamus.

## Introduction

It is now well-established that early life pain and stress have both acute and lasting consequences on brain development and affective behaviors, stress responsiveness, and pain thresholds in both clinical populations and pre-clinical animal models [1–11]. One source of early life adversity is medical trauma that occurs in the neonatal intensive care unit (NICU) in premature or sick neonates. While increases in NICU admissions have coincided with lower levels of infant mortality [12–14], the pain and stress that occur in those settings has been linked with later life dysfunction including mental health vulnerability [6,7,15–19]. However, the mechanisms underlying these changes are still being elucidated, which is a critical impediment to the proper prevention, diagnosis, and treatment of neonates who are exposed to early life medical trauma.

Early life events have been well established to have lasting impacts on developmental trajectory of the brain and nervous system, despite the existence of infantile amnesia and stress hyporesponsive periods [20–26]. While neonates must continue to stay close to sources of food and protection despite any painful or adverse stimulation [27], plasticity during critical periods of development is necessary to prepare organisms for the particular environmental challenges they will face. Thus, early life adversity can serve to prepare organisms for stressful or painful futures by priming either resilience or vulnerability, which then can manifest during later-life stress or trauma (often called a “two-hit model” when discussing mental health outcomes) [28–31]. Indeed, pre-clinical models have demonstrated several effects of early life pain and stress. For example, brief handling and maternal separation paradigms have also been commonly used to examine the neurobiology of early life stress, which has demonstrated changes in cortisol/corticosterone expression and activation of a variety of stress-and emotion-relevant brain structures [32–40]. In particular, early handling (consisting of repeated short periods of separation and experimenter handling) differs from maternal separation (characterized by long periods of separation of the dam and pups, typically without explicit experimenter handling) in the direction of the effect on the HPA-axis and cortisol expression, with early handling leading to reduced stress responsiveness and maternal separation leading to elevated stress responsiveness, although all of these effects are highly parameter dependent [37,40,41].

The NICU involves acute and repeated pain, in addition to non-painful stress, which may cause different or additional effects. For example, a NICU-like “pup in a tea-ball” model with paw pricks produces changes in the hypothalamus-pituitary-adrenal (HPA-)axis responsiveness in females [42] and changes to glutamate/GABA signaling in the pre-frontal cortex and hippocampus [43]. Our lab, along with others, has shown sex-dependent changes in corticotropin releasing factor (CRF) signaling in the amygdala due to handling [44], cold stress [26,45], and pin-prick pain [46,47]. While CRF (also called CRH) in the amygdala has been increasingly implicated in pain [48,49], it was first identified as a key mechanism of hypothalamic control of the HPA-axis with significant roles in early life stress and sex-specific changes [50,51]. Overall, the effects of early life adversity in the amygdala, as well as the hypothalamus and HPA-axis, may be of critical importance, as these structures are involved in threat, stress, and pain processing, as changes in forebrain CRF levels during development have been linked with long-term changes in affective and sensory function [21].

Importantly, CRF signaling in the brain can be altered not only by changes in expression of the CRF ligand, but also by changes in the CRF receptor. CRF has two major receptors, the high affinity CRFR1 and the lower affinity CRFR2. These receptors are often characterized as having generally opposing functions in stress, fear and pain, with CRFR1 activation leading to enhancement of stress, fear and pain and CRFR2 activation leading to recovery, diminished stress, fear and responding [52–56], although this simplistic view is likely brain region and context dependent [57]. Changes in the relative expression of CRF receptor 1 and receptor 2 have been linked to chronic pain conditions [52,58] and early life stress [38,59,60], including early life inflammatory pain [61].

This paper follows up on prior work by examining the effects of an early life pain model, designed to capture key features of the NICU, on CRF receptor mRNA expression. Examining the neurobiological consequences of NICU-like experiences is a critical topic of study. Many sources of early life pain and stress are unavoidable and likely lead to similar neurobiological consequences. However, the NICU represents an environment in which there is both high motivation and ability to enact positive changes to improve later outcomes. Our hypothesis involves a two-hit model, in which we believe that early life events prime the nervous system to respond in an altered manner to a later life stressor or trauma. In our case, the initial trauma is designed to model, albeit incompletely, the NICU experience, with repeated removal from the dam, handling, and skin-breaking paw pricks over the first week of life. Then, later in life, a traumatic event in the form of classical fear conditioning activates a susceptible amygdala and hypothalamus, leading to changes in pain thresholds and affective expression, with both vulnerable and resilience phenotypes emerging, depending on the parameters [31,46,62,63]. We have previously found sex-dependent changes in the number of CRF-expressing cells in the amygdala (with an acute increase, followed by a lasting decrease) and to a lesser extent, the hypothalamus [46,47]. The current experiments examine whether our early life pain model induces changes in the expression of CRF receptors within the amygdala and hypothalamus.

## Methods

### Subjects

Male and female Sprague Dawley rats were bred in-house using a protocol previously described [31]. We used a total of 89 pups (42m and 47F) from 16 (8 NICU-treated; 8 undisturbed) litters. All litters were housed in (W x D x H) 39.5 cm x 34.6 cm x 21.3 cm closed-environment cages (Tecniplast, West Chester PA). On postnatal day (PD) 1, pups were removed from their mother, placed on a heating pad, sexed, marked via crystal violet stain, and culled to no more than 10 rats per litter (5 males and 5 females when possible). Pups were weaned on PD 21 and lived with their same-sex littermates (approximately 5 per cage). No more than one same sex littermate was assigned to each experimental group, with experimental group defined as a combination of sex (male or female), neonatal condition (early pain, handled, undisturbed), and fear conditioning status (fear conditioned or not). All rats were maintained on a 12:12 light/dark cycle with lights on at 07:00. Food and water were available ad libitum, and at the end of experimentation, rats were euthanized via pentobarbital overdose and brains were collected. All rats were treated in accordance with the NIH *Guide for the Care and Use of Laboratory Animals* (2011) and all procedures were approved by the University of New England’s Institutional Animal Care and Use Committee (IACUC). See Figure 1 for a timeline of procedures.

**Figure 1.**
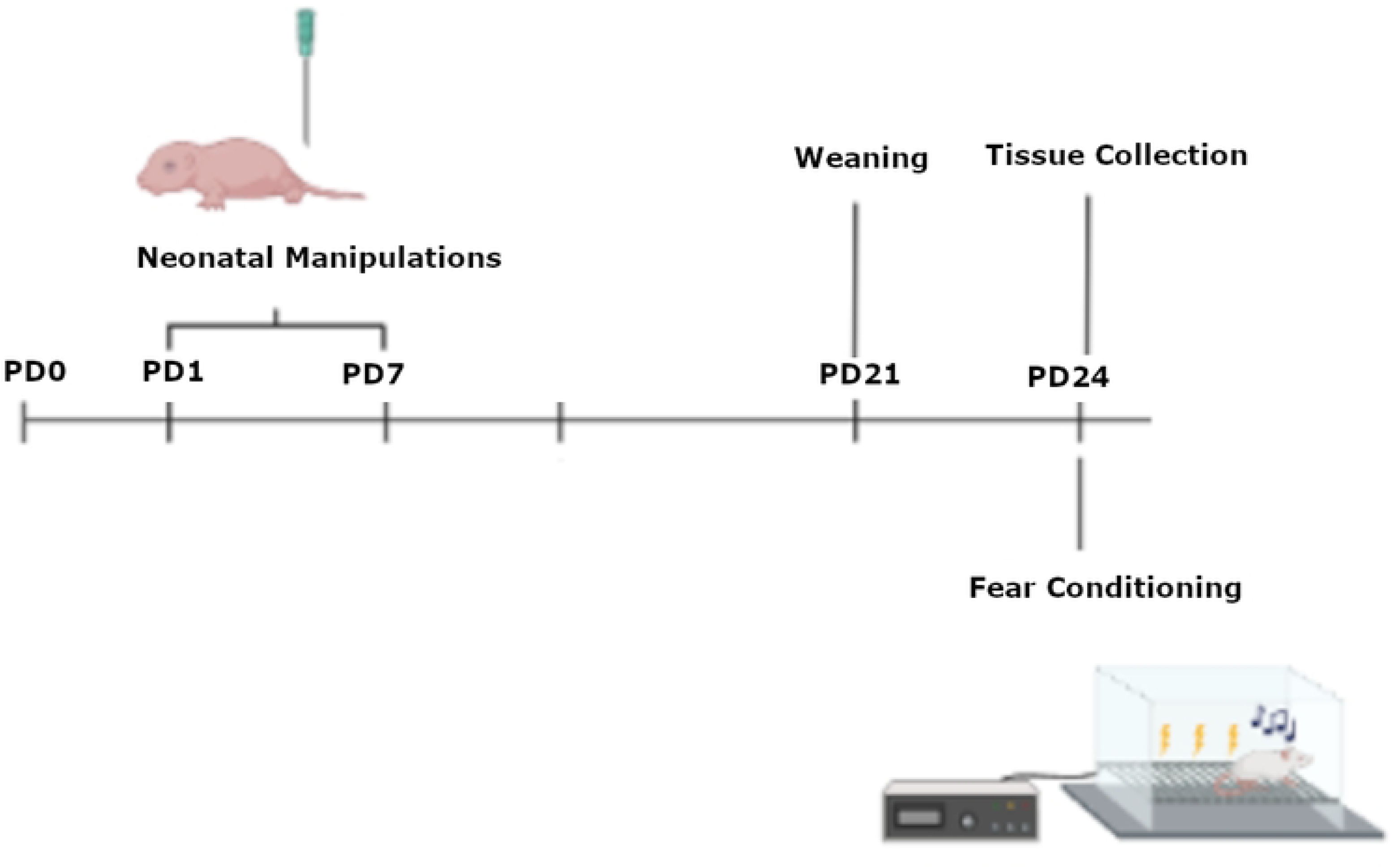
A timeline of procedures. Rats were born on PD0, were subjected to neonatal manipulations from PD 1 - 7, weaned on PD 21 and fear conditioning on PD 24 prior to tissue collection.

### Neonatal Pain

The procedure was similar to that previously described (S. M. Davis et al., 2018, 2021). Briefly, for early pain litters, on PDs 1-7, rats were removed from the dam, placed on a heating pad and received either a left hindpaw prick using a 24 G needle (NICU-treated) or a non-painful tactile touch (handled) every two hours, four times per day starting at 09:00 am. The needle prick procedure is limited to breaking the dermal layer of the hind paw and eliciting a drop of blood. Non-painful touch using the tip of the experimenter’s finger in the same location that was pricked in the early pain-treated subjects. For each of the 4 daily pricking events, subjects were separated from the dam for approximately 5-10 min, with 5 min being the minimum separation.

During this period, subjects were repeatedly handled by the experimenter while separated from each other. Pups were reunited shortly prior to being returned to the dam. Subjects were marked with crystal violet periodically (typically on PD 1 and 4) to identify individual pups. Control litters were left undisturbed.

### Apparatus

Fear conditioning was performed in four Startfear chambers (Harvard Apparatus/Panlab model #58722) with two separate contextual cues that differed in shape (square vs. circle), color (black vs. white walls), and scent (70% ethanol vs. 1% ammonia).

### Fear Conditioning

On PD 24 fear conditioning rats underwent a fear conditioning protocol, similar to that previously described [31,64]. Rats were placed in their preassigned (counterbalanced) fear conditioning chamber and the program was initiated. The percent of time spent freezing during the first 5 mins was recorded (Habituation). Following the habituation period, a 67-dB tone conditioned stimulus (CS) was presented for 10 sec, immediately followed by a 2-s 1.0-mA foot shock serving as the unconditioned stimulus (UCS). There were 10 tone-shock pairings, separated by an average inter-trial interval (ITI) of 2.5 minutes. Subjects were euthanized approximately 30 minutes after this procedure (∼1 hr after onset of conditioning) and tissue was collected.

### Tissue Collection

Subjects were euthanized with a volume of 0.25ml pentobarbital (390mg/ml), prior to intra-cardiac perfusion of saline, followed by 4% paraformaldehyde (PFA) on PD 24. Brains were collected and subsequently post fixed for 24 hours in 4% PFA. All animal euthanasia was consistent with the American Veterinary Medical Association procedures. After the 24-hour post-fix, brains were cryoprotected in 30% sucrose. When cryoprotection was finished, brains were embedded in Tissue-Tek O.C.T. Compound (Sakura Finetek) and flash frozen using liquid nitrogen. Brains were then stored at-80C prior to cryosectioning at 15um onto Superfrost plus microscope slides. Emphasis was placed to identify two sections between-1.8 and-2.4 mm from bregma for consistency between subjects and optimal amygdala presentation. These sections were stored at-80C until application of our RNAscope® Fluorescence in situ hybridization protocol (approximately 1 month).

### Fluorescence in situ hybridization

All fluorescence in situ hybridization (FISH) was performed using a commercially available system [RNAscope; Advanced Cell Diagnostics (ACD)] and utilized probes targeting *Crhr1* (product number: 318911) and *Crhr2* (product number: 417851-C2). Our protocol was developed using RNAscope Multiplex Fluorescent Reagent Kit v2 user manual (document number: 323100-USM), manufacturer technical note regarding tissue detachment, manufacturer modifications for fixed frozen tissue (ACD), and our previous work [47]. In the concluding steps of our protocol, DAPI was applied to brain sections as a counterstain for cellular and region identification.

Sections were kept at-80 until staining. For target retrieval, sections were warmed for 1 hour at room temperature, then dried at 60°C for 45 minutes, prior to a post fix in 4%PFA for 1 hour. They were washed in ascending concentrations of ethanol (50%, 70% and 100%), treated with hydrogen peroxide, washed in DI water, and placed in the target retrieval solution for 15 minutes in a steamer at 99°C, prior to a wash in DI water followed by 100% ethanol. Sections were then dried at 60°C for 30 minutes prior to being left overnight. *In situ hybridization* began with application of the probes and baking at 40°C for 2 hours. Each channel was then hybridized with the appropriate AMP solution for 30 minutes at 40°C. The appropriate HPR was then added at 40°C for 15 min, prior to the application of the dye. Washing using the wash buffer occurred between each step.

RNAscope assays were conducted in batches. We identified two neighboring sections from each subject that appeared to contain the subject regions. Each RNAscope batch contained one section from each condition (sex x neonatal condition x juvenile treatment) from tissue collected on the same day. Additionally, RNAscope batches were duplicated such that the two neighboring sections from the same brain were stained in separate batches to control for any inter-batch variability. Lastly, subject averages (for each hemisphere) were created from the two sections and these averages were used for all subsequent analysis. Thus, each subject contributed only one value (the subject average) per hemisphere to any statistical analysis. Per manufacturer instructions (RNAscope; ACD) to ensure consistency between subsequent FISH batches, each batch consisted of additional sections processed with either positive control probe [product number: 320891; containing Polr2a (channel 1), PPIB (channel 2)] or negative control probe [product number: 320871; containing DapB gene accession EF191515 from the SMY strain of Bacillus subtilis (channels 1 and 2)].

Fluorescence multiplex imaging was done between 2 days and 2 weeks after FISH. All quantified images and channels were taken under the same magnification (20x) and exposure settings (DAPI – 8ms; FITC channel (*crhr1*) 300ms; CY3 channel (*crhr2*) – 35ms) on an Olympus VS200 slide scanner. Images were analyzed using the FIJI package of NIH’s open-source image analysis software ImageJ [65,66]. Image analysis and region of interest (ROI) identification occurred using encoded brain numbers keeping the experimenter blind to the identity of the image being analyzed.

Images were first subject to channel separation and the DAPI channel was used to Identify regions of quantification (Central Nucleus of the Amygdala; CeA, Basolateral Amygdala; BLA, Periventricular Nucleus; PVN and Ventromedial Hypothalamus; VMH) (See Figure 2). The DAPI channel was then dilated and used to create a cell mask for these regions. Due to the different levels of background and positive staining, different thresholding algorithms were used on each channel. FIJI’s Triangle Theshold was used for the green channel and the RenyEntropy Threshold for the red channel. The number of cells containing *crhr*1 and *crhr2* was assessed, as well as the overall area of the region of interest displaying positive signaling. Additionally, luminance of straining was assessed in these regions via the averaging of grayscale values, as a control. There were no differences in luminance of staining found, suggesting consistent staining and imaging quality across all groups. All images were subject to the same thresholding, processing, and quantification methods.

**Figure 2:**
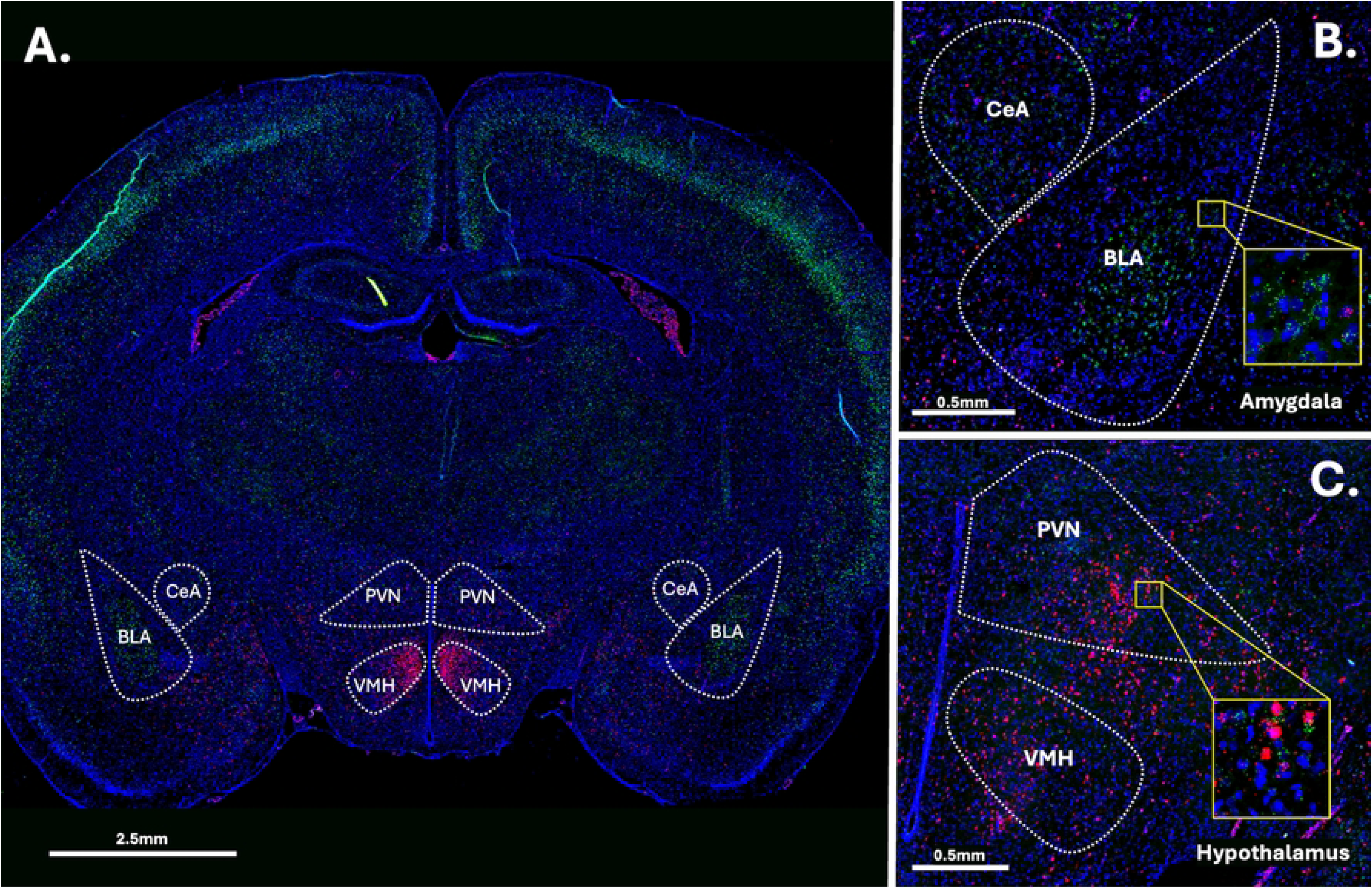
Example image of RNAscope straining. Panel A: Full brain section straining. Panel B: Closeup on the amygdala. Panel C. Closeup on the hypothalamus. Green = crfr1, Red = crfr2. Blue = DAPI.

### Experimental Design and Analysis

Data were analyzed using a 2 (sex) X 3 (Neonatal condition: early pain, handled, undisturbed) X 2 (juvenile treatment: fear conditioning or not) X 2 (hemisphere – a repeated measure) mixed model GLM with IBM SPSS version 27. No more than one same sex littermate was assigned to each experimental group. Individual data points were removed as statistical outliers if they were identified as extreme outliers using SPSS’s outlier analysis in their respective group. No more than 1 data point per group (combination of sex, neonatal condition, juvenile condition and hemisphere) met this criterion with 5 total data points in the BLA, 2 in the CeA, 6 in the PVN and 3 in the VHM across all measures. In addition, sections were removed if the region of interest was not present, was damaged, or was otherwise not quantifiable.

3 measures are reported for each target, the number of cells expressing that target, the percentage of DAPI-stained cells that express the target and the overall percentage of pixels in the ROI that expressed the signal. Examining differences in the number of cells tests the hypothesis that there is a recruitment or decline in the population of cells that express the CRF receptors, whereas examining the percentage of pixels expressing a target is also sensitive to changes in the degree of expression within the existing cell populations. Data are reported as mean ± SEM and a *p*-value of ≤ 0.05 was considered statistically significant, while *p*-values between 0.05 and 0.10 were considered trends towards significance. A Tukey’s post hoc analysis was performed when there were more than two levels of a variable to compare. In addition to graphing the overall data, statistically significant differences were highlighted with additional figures.

## Results

### Basolateral Amygdala

After quality control and outlier analysis, between 82-84 brains contributed to each measure. There was a minimum of 5 and a maximum of 8 brains included per group.

Number of nuclei stained with DAPI (Figure 3A): These data were first subjected to a 2 (Sex: M,F) X 3 (Neonatal condition: undisturbed, handled, early pain) X 2 (Juvenile treatment: Fear Conditioned or not) X 2 (hemisphere – a repeated measure) mixed model General Linear Model (GLM) for the number of nuclei stained with DAPI. Surprisingly, we found a significant effect of juvenile treatment F(1,70) = 5.058, p<.05 η^2^ =.07 (Figure 3B) with fear conditioned subjects showing more expression and a significant effect of hemisphere F(1,70) = 32.377, p<.01 η^2^ =.316 (Figure 3C) with the left side showing more expression. Further analysis demonstrated that these differences were attributable to incidental differences in ROI volume, which moderately correlated (*r*s =.40-.60 for each group) with the number of DAPI-stained nuclei and showed the same pattern of statistical differences. There were no differences in nuclei density. Given that ROIs were created blind to condition, we believe that these differences were spurious, but should be considered when interpreting later results. No other significant differences were found in this measure. For a list of all analyses, see Table 1.

**Figure 3:**
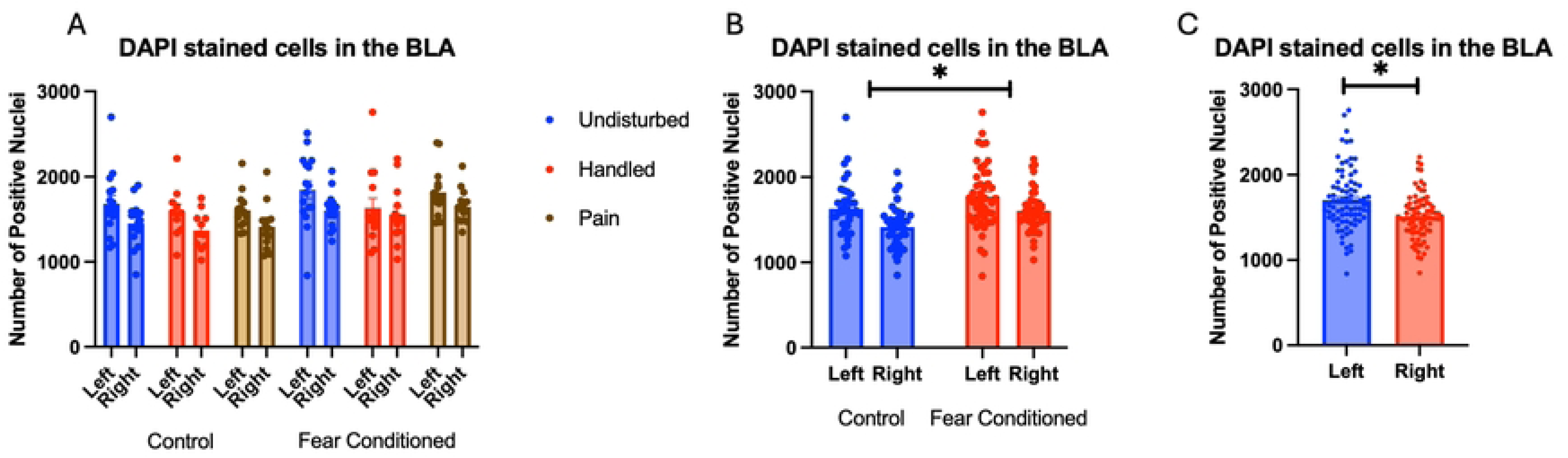
DAPI stained nuclei in the BLA. PD 1-7 male and female rats were subjected to a painful neonatal experience, non-painful removal from the dam and handling, or left undisturbed except for normal animal colony procedures. Subjects were either fear conditioned or not on PD 24. Tissue was collected and subjected to RNAscope in situ hybridization in both hemisphere as shown in Panel A. There were significantly more DAPI-stained nuclei in fear conditioned subjects (Panel B) and in the left hemisphere (Panel C). These differences were attributable to incidental differences in ROI size.

**Table 1:**
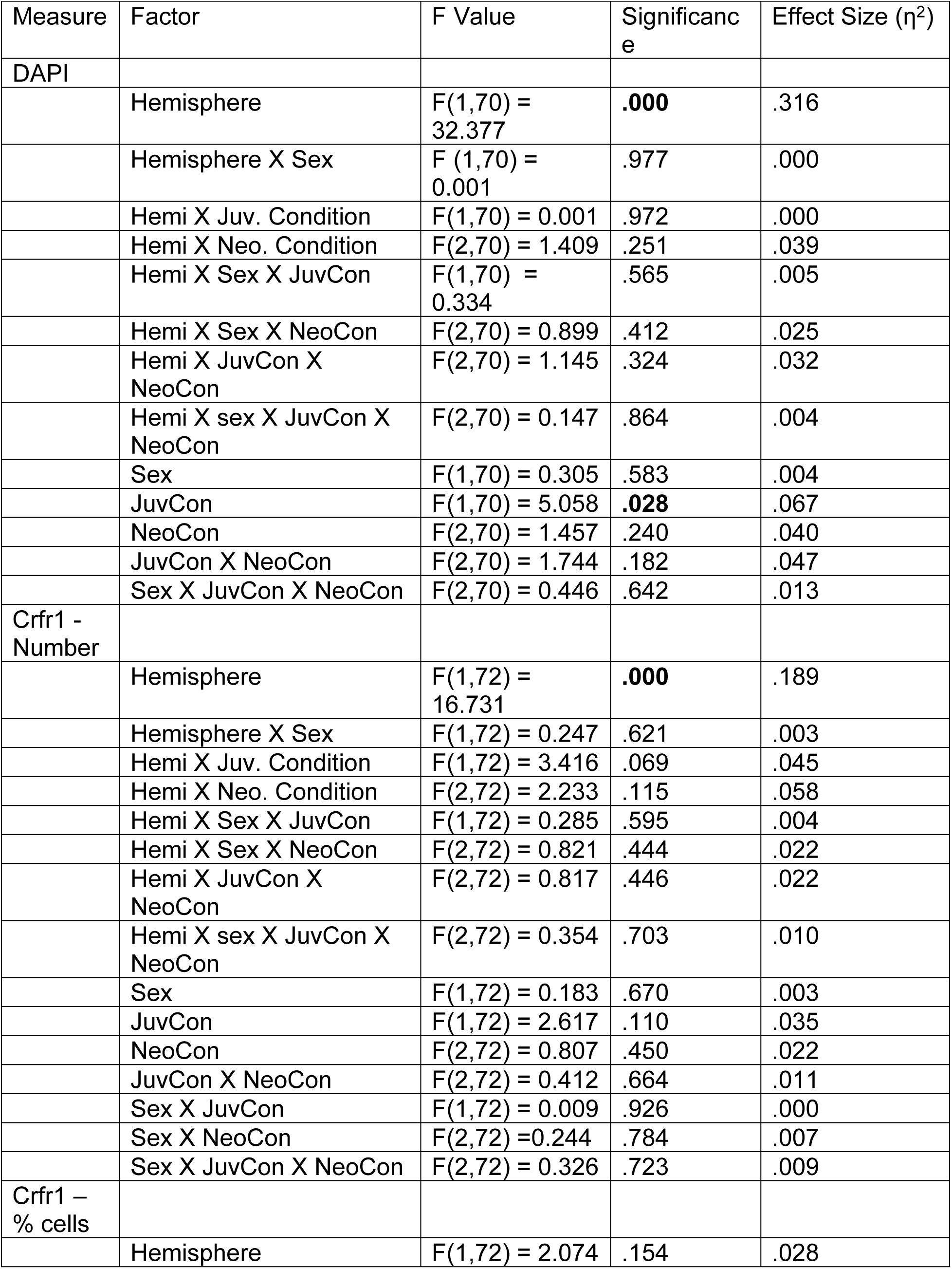
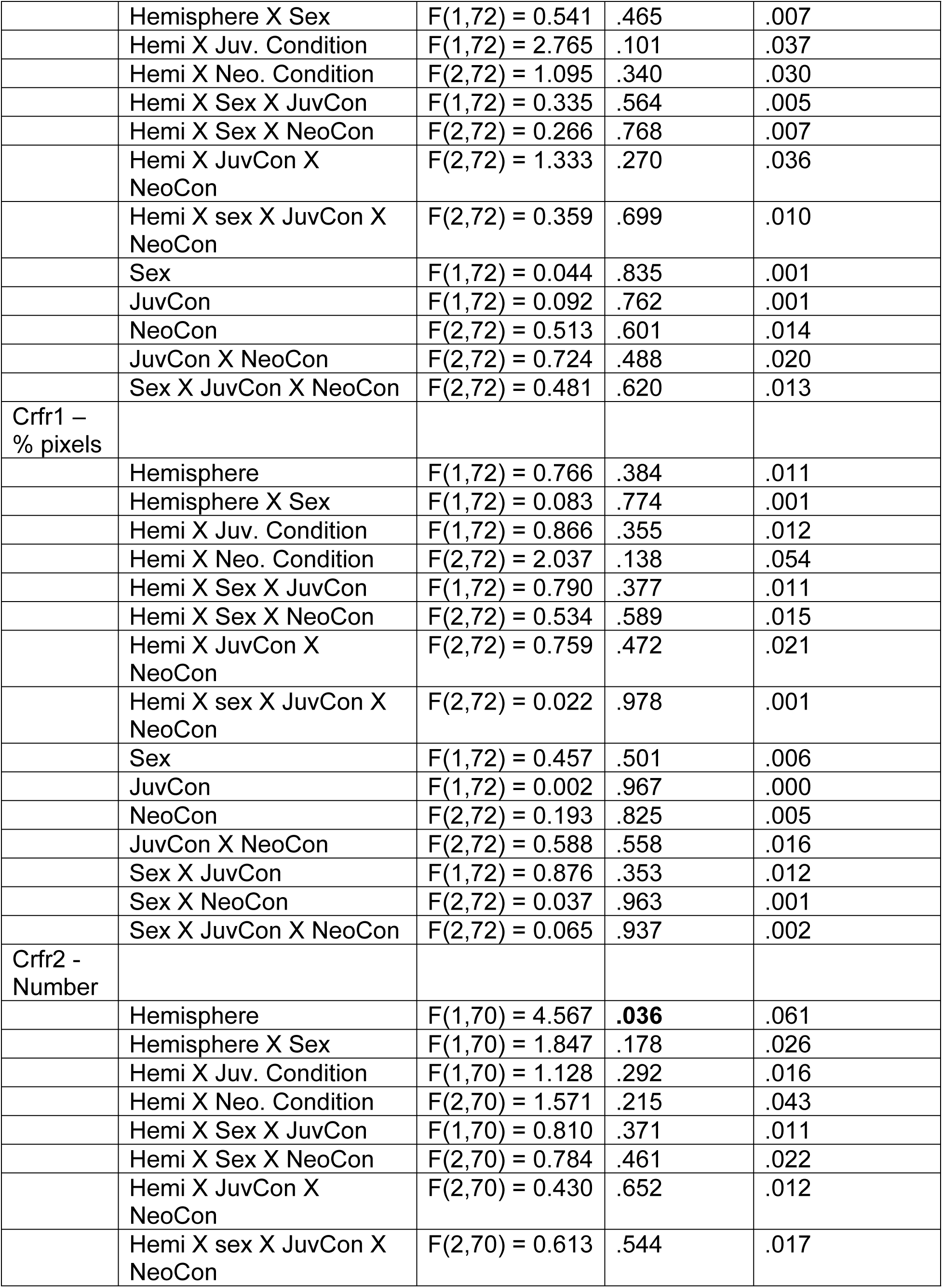
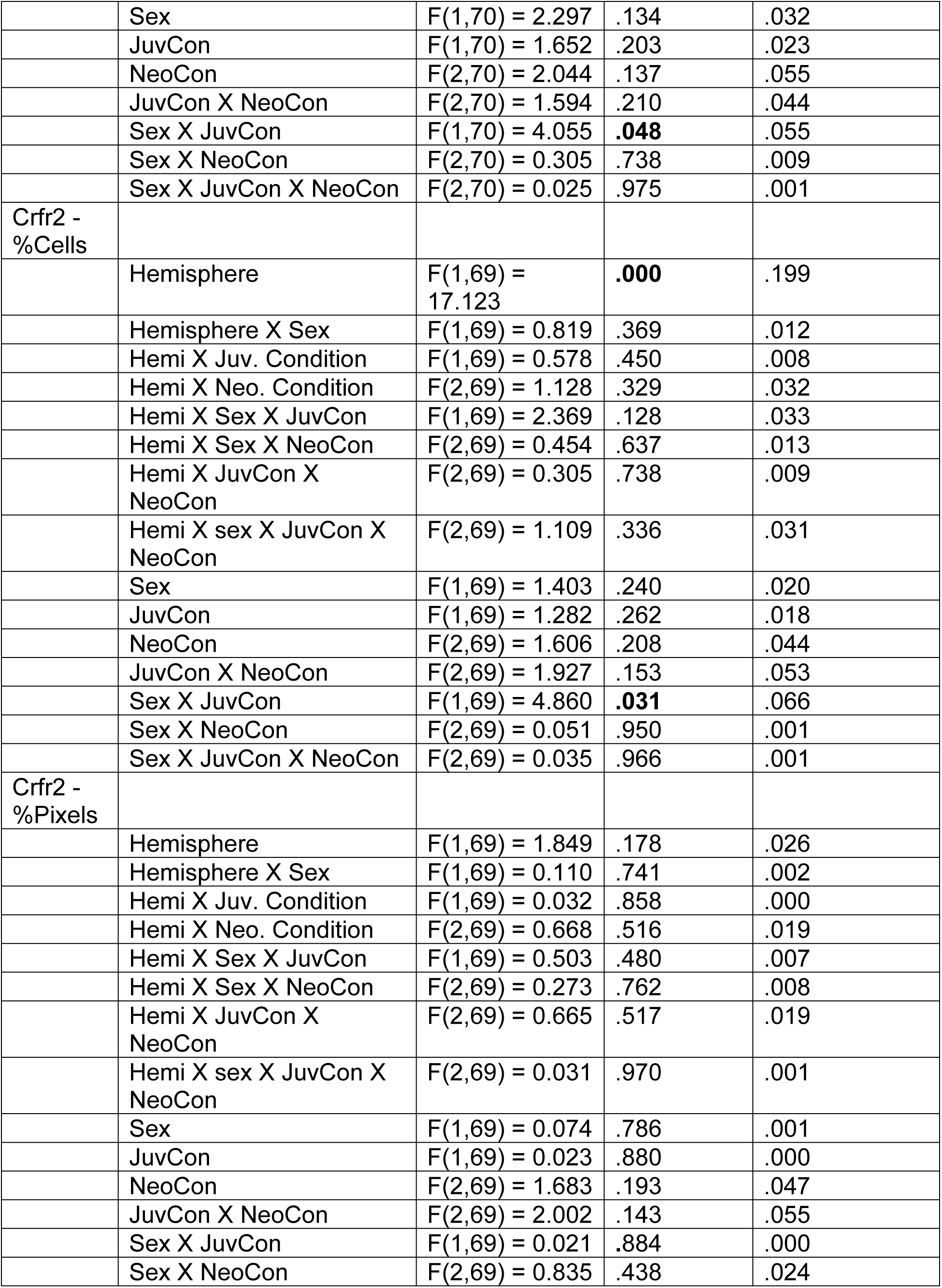
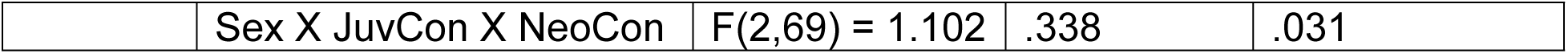
BLA.

Number of Cells expressing *crhr1*: These data were first subjected to a 2 (Sex: M,F) X 3 (Neonatal condition: undisturbed, handled, early pain) X 2 (Juvenile treatment: Fear Conditioned or not) X 2 (hemisphere) mixed model General Linear Model (GLM) for the number of cells expressing *crhr1.* There were no significant main effects or interactions with sex, so we combined across sex (Figure 4A). We observed a significant main effect of hemisphere F(1,72) = 16.73, p<.01 η^2^ =.189, and a trend towards a significant hemisphere X juvenile treatment interaction F(1,72) = 3.62, p<.07 η^2^ =.045. A follow-up analysis demonstrated a significant effect of hemisphere only in the fear conditioned subjects F (1,38) = 18.101, p<.01 η^2^ =.323 and not in control subjects F (1,34) = 2.458, n.s. η^2^ =.067 with the left hemisphere demonstrating significantly more expression (*Figure 4B*).

**Figure 4:**
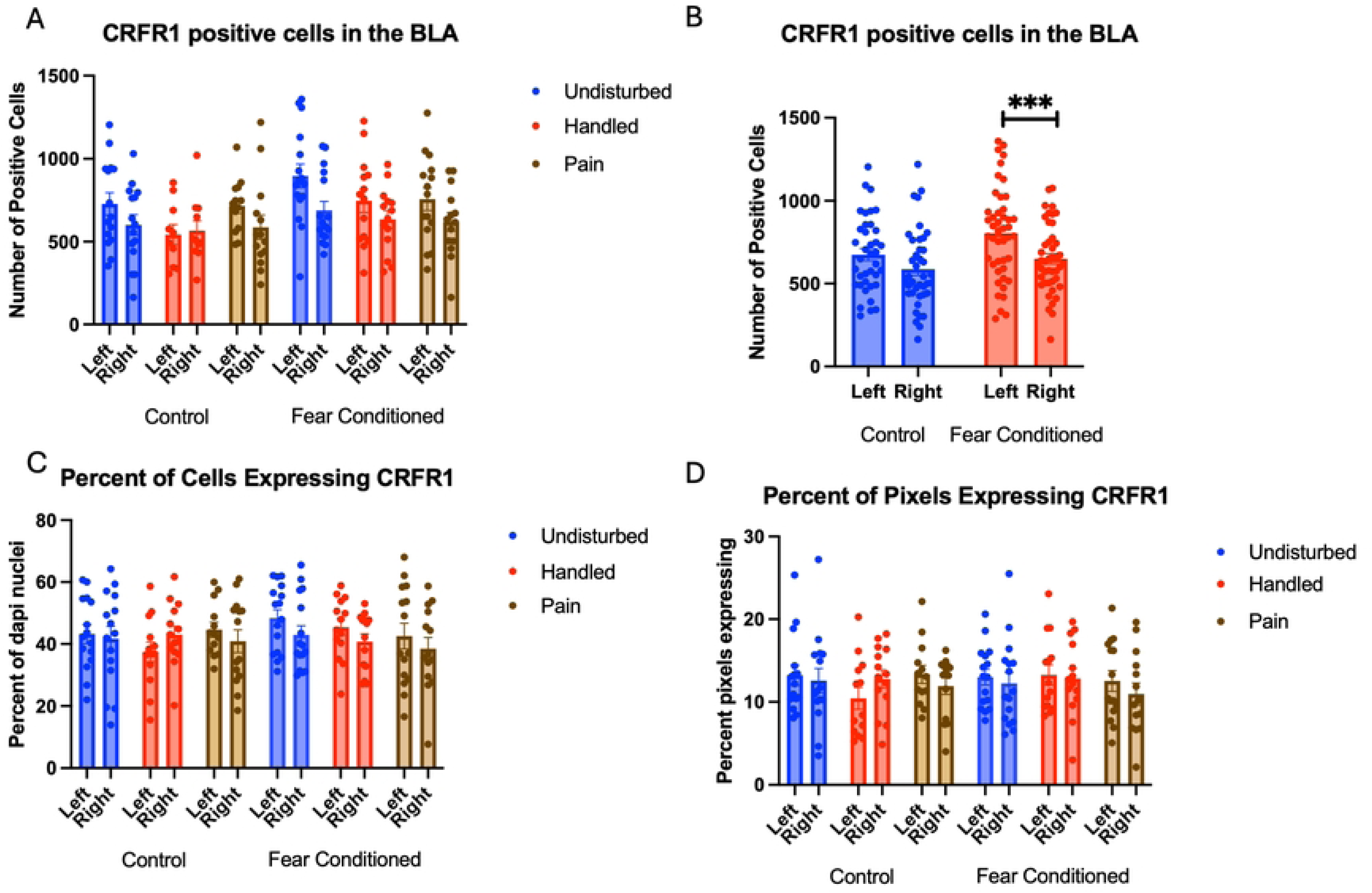
Crhr1 expression in the BLA. PD 1-7 male and female rats were subjected to a painful neonatal experience, non-painful removal from the dam and handling, or left undisturbed except for normal animal colony procedures. Subjects were either fear conditioned or not on PD 24. Tissue was collected and subjected to RNAscope in situ hybridization in both hemisphere. Panel A: All data from both hemispheres, collapsed across sex due to lack of statistical differences. Panel B: Data collapsed across all variables except juvenile treatment and hemisphere, to highlight the interaction. *** = statistically significant p<.001. Panel C: the percent of DAPI stained nuclei that express crhr1. Panel D: the percent of pixels that express crfr1.

A 2 (Sex: M,F) X 3 (Neonatal condition: undisturbed, handled, early pain) X 2 (Juvenile treatment: Fear Conditioned or not) X 2 (hemisphere) mixed model MANOVA examining percent of cells expressing *crhr1* revealed no significant differences (Figure 4C), as did a similar analysis on percent of pixels (Figure 4D). As the number, but not the percent of cells or pixels differ, the hemispheric difference in the number of cells expressing *crfr1* is likely an artifact reflecting differences in the number of DAPI nuclei. All statistics can be found in Table 1.

Number of Cells expressing *crhr2*: These data were also first subjected to a 2 (Sex: M,F) X 3 (Neonatal condition: undisturbed, handled, early pain) X 2 (Juvenile treatment: Fear Conditioned or not) X 2 (hemisphere) mixed model GLM for the number of cells expressing *crhr2* in the BLA (Figure 5A). We observed a significant main effect of hemisphere F(1,70) = 4.567, p<.05 η^2^ =.061, with the right hemisphere demonstrating higher levels of expression and a significant sex X juvenile treatment interaction F(1,70) = 4.06, p <.05 η^2^ =.055, which demonstrated significantly more *crhr2* expression in the right hemisphere of fear conditioned subjects only in females, with a moderate effect size (Figures 5B and 5C). Importantly, this cannot be accounted for by the higher levels of DAPI expression in the left hemisphere.

**Figure 5:**
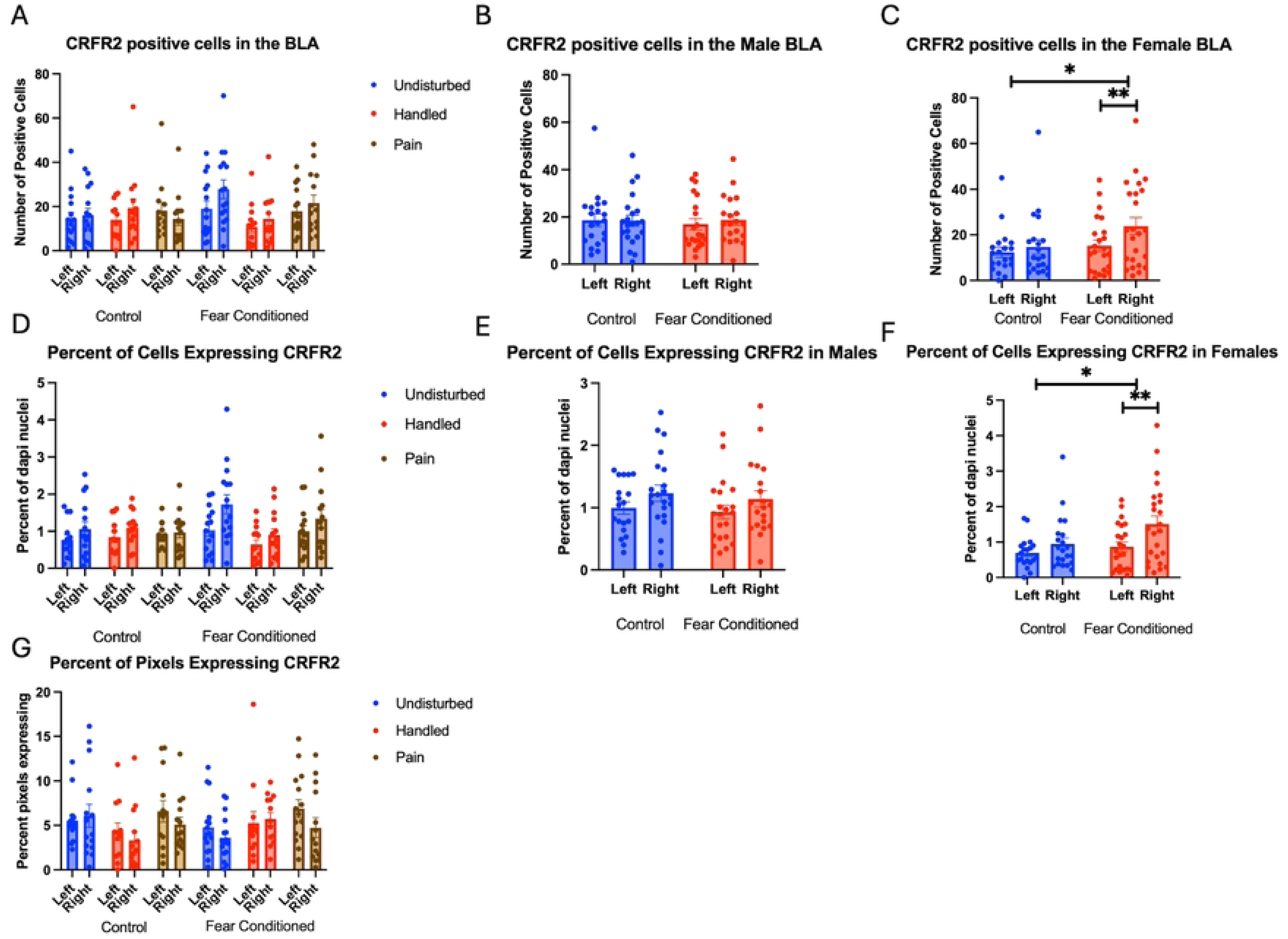
Crhr2 expression in the BLA. PD 1-7 male and female rats were subjected to a painful neonatal experience, non-painful maternal removal and handling, or left undisturbed except for normal animal colony maintenance. Subjects were either fear conditioned or not on PD 24. Tissue was collected and subjected to RNAscope in situ hybridization in both hemisphere. Panel A: The number of cells expressing crhr2 from both hemispheres, collapsed across sex. Panel B: The number of cells expressing crhr2 from male subjects highlighting the lack of effect of fear conditioning. Panel C: The number of cells expressing crhr2 from female subjects highlighting the effect of fear conditioning and hemisphere. * = statistically significant p<.05. Panel D: Percentage of cells expressing crhr2 collapsed across sex. Panel E: Percentage of cells expressing crhr2 highlighting the lack of effect of fear conditioning and hemisphere. Panel F:: The percentage of cells expressing crhr2 from female subjects highlighting the effect of fear conditioning and hemisphere * = statistically significant p<.05. Panel G: The percentage of overall ROI expressing positive signal after thresholding.

Percent of DAPI cells that express *crhr2*: These data were also first subjected to a 2 (Sex: M,F) X 3 (Neonatal condition: undisturbed, handled, early pain) X 2 (Juvenile treatment: Fear Conditioned or not) X 2 (hemisphere) mixed model GLM for the number of cells expressing *crhr2* in the BLA (Figure 5D). We again observed a significant main effect of hemisphere F(1,69) = 17.123, p<.01) η^2^ =.199, with the right hemisphere demonstrating higher levels of expression (and a large effect size) as well as a significant sex X juvenile treatment interaction F(1,69) = 4.860, p <.05 η^2^ =.066, which demonstrated significantly more *crhr2* expression in the right hemisphere of fear conditioned subjects only in females (Figures 5E and 5F). Because this is a percentage and opposite to the hemisphere with more DAPI, this difference also cannot be accounted for by any difference in the number of cells examined.

A 2 (Sex: M,F) X 3 (Neonatal condition: undisturbed, handled, early pain) X 2 (Juvenile treatment: Fear Conditioned or not) X 2 (hemisphere) mixed model GLM for the percent of pixels expressing *crhr2* in the BLA found no significant differences (Figure 5G). All statistics can be found in Table 1.

### Central Nucleus of the Amygdala

A total of 83-87 brains survived quality control and outlier analysis for the CeA, for each measure. There was a minimum of 6 and a maximum of 8 subjects per group.

Number of nuclei stained with DAPI (Figure 6A). These data were again subjected to a 2 (Sex: M,F) X 3 (Neonatal condition: undisturbed, handled, early pain) X 2 (Juvenile treatment: Fear Conditioned or not) X 2 (hemisphere) mixed model GLM for the number of stained nuclei. We once again found an unexpected significant effect of juvenile treatment (Figure 6B) F(1,75) = 8.395, p<.01 that correlated with differences in ROI area and reflected more cells in fear conditioned subjects. This is also likely to be spurious, but subsequent analyses will be considered in light of this difference.

**Figure 6:**
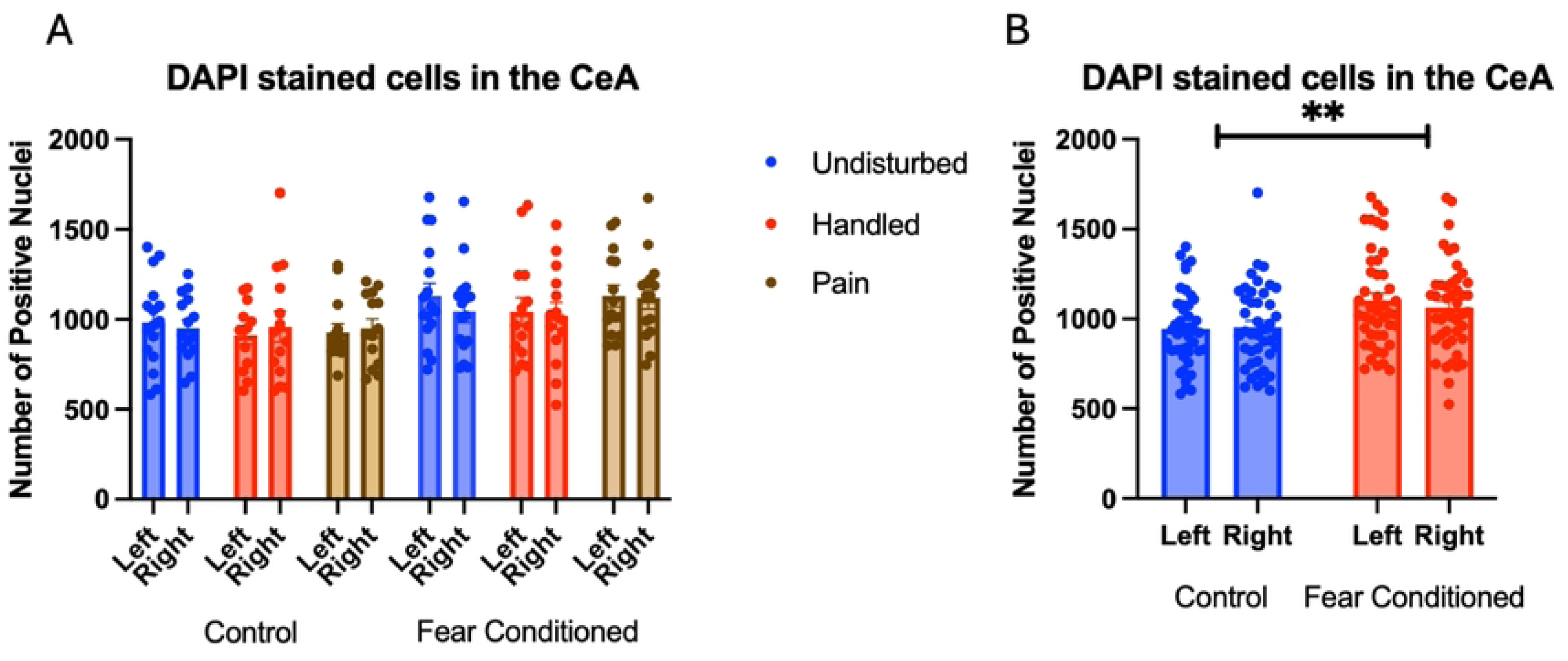
DAPI stained nuclei in the CeA. PD 1-7 male and female rats were subjected to a painful neonatal experience, non-painful removal from the the dam and handling, or left undisturbed except for normal animal colony procedures. Subjects were either fear conditioned or not on PD 24. Tissue was collected and subjected to RNAscope in situ hybridization in both hemisphere as shown in Panel A. There were significantly more DAPI-stained nuclei in fear conditioned subjects (Panel B). These differences were attributable to incidental differences in ROI size.

Number of Cells expressing *crhr1.* Similar to above, these data were first subjected to a 2 (Sex: M,F) X 3 (Neonatal condition: undisturbed, handled, early pain) X 2 (Juvenile treatment: Fear Conditioned or not) X 2 (hemisphere) mixed model GLM for the number of cells expressing *crhr1.* With no sex differences, data are collapsed across this variable *(Figure 7A).* The initial analysis found a trend toward a significant hemisphere X juvenile treatment interaction F(1, 74) = 3.83, p=.054 η^2^ =.049 (Figure 7B). A follow up analysis showed a trend towards greater expression in left hemisphere of fear conditioned subjects compared to controls (p=.065; η^2^ =.045). This can be accounted for by the increase in DAPI expression by fear conditioned subjects and likely does not reflect an actual effect of conditioning. A 2 (Sex: M,F) X 3 (Neonatal condition: undisturbed, handled, early pain) X 2 (Juvenile treatment: Fear Conditioned or not) X 2 (hemisphere) mixed model MANOVA examining percent of cells expressing *crhr1* revealed no significant differences (*Figure 7C*), as did a similar analysis on percent of pixels (Figure 7D). Thus, similar to the BLA, the difference in the number of cells expressing *crfr1* is likely an artifact reflecting differences in ROI area altering the number of DAPI nuclei. All statistics can be found in Table 2.

**Figure 7:**
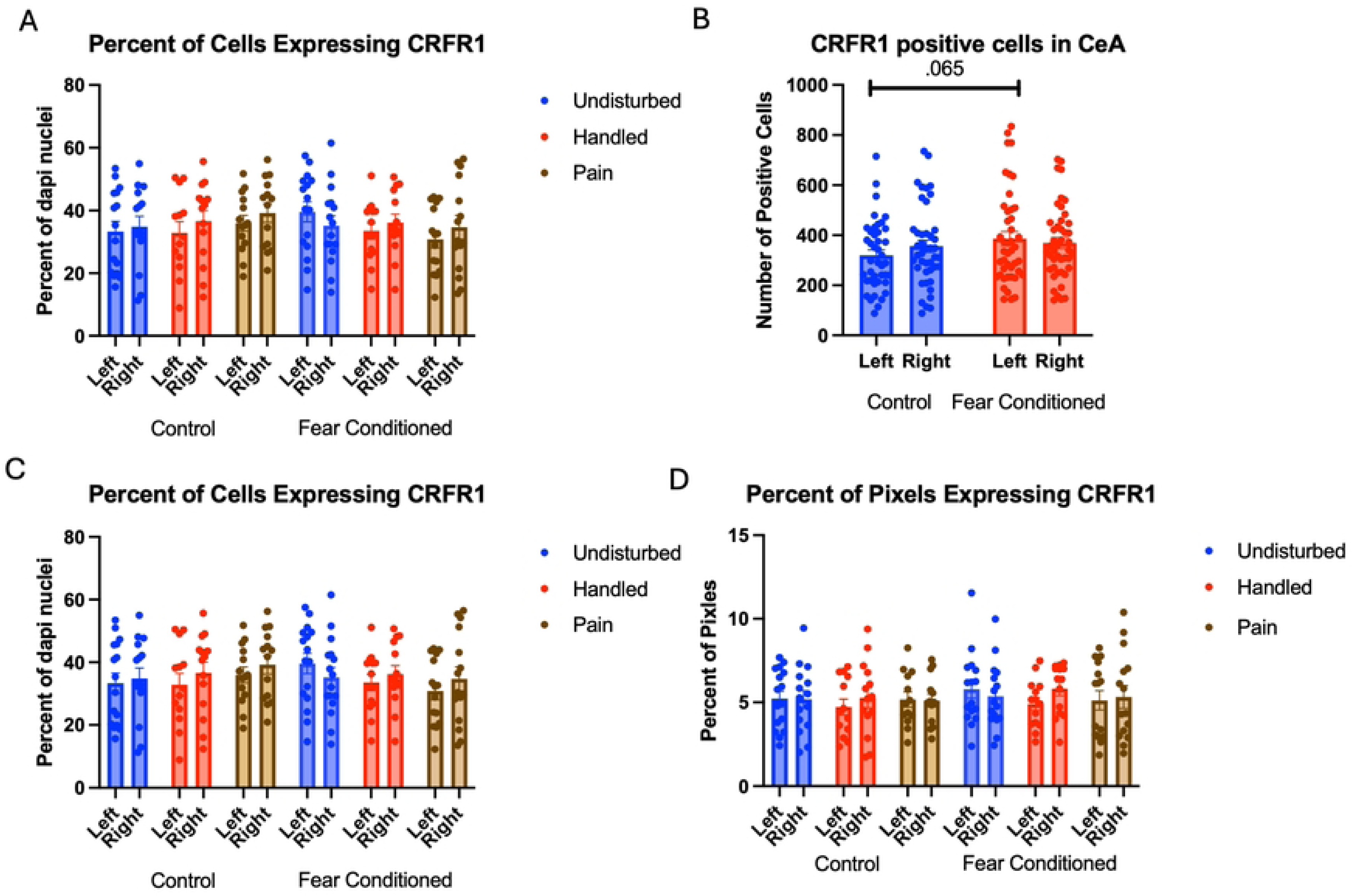
crhr1 expression in the CeA. PD 1-7 male and female rats were subjected to a painful experience, non-painful removal from the dam and handling, or left undisturbed except for normal animal colony maintenance. Subjects were either fear conditioned or not on PD 24. Tissue was collected and subjected to RNAscope in situ hybridization in both hemispheres. Panel A: Number of cells expressing crhr1 from both hemispheres, collapsed across sex. Panel B: Number of cells expressing crhr1 collapsed across neonatal condition and sex, to highlight the hemisphere by juvenile condition statistical interaction. Panel C: Percentage of cells in overall ROI expressing crhr1. Panel D: Percentage of pixels expressing positive signal after thresholding. There were no significant differences between groups.

**Table 2:** CeA.

| Table 2: CeA |  |  |  |  |
| --- | --- | --- | --- | --- |
| Measure | Factor | F Value | Significance | Effect Size ($\eta^2$) |
| DAPI |  |  |  |  |
|  | Hemisphere | F(1,75) = 0.130 | .719 | .002 |
|  | Hemisphere X Sex | F(1,75) = 0.131 | .719 | .002 |
|  | Hemi X Juv. Condition | F(1,75) = 1.884 | .174 | .025 |
|  | Hemi X Neo. Condition | F(2,75) = 0.839 | .436 | .022 |
|  | Hemi X Sex X JuvCon | F(1,75) = 0.576 | .450 | .008 |
|  | Hemi X Sex X NeoCon | F(2,75) = 0.749 | .476 | .020 |
|  | Hemi X JuvCon X NeoCon | F(2,75) = 0.137 | .872 | .004 |
|  | Hemi X sex X JuvCon X NeoCon | F(2,75) = 2.631 | .079 | .066 |
|  | Sex | F(1,75) = 0.779 | .380 | .010 |
|  | JuvCon | F(1,75) = 8.395 | **.005** | .101 |
|  | NeoCon | F(2,75) = 0.328 | .721 | .009 |
|  | Sex X JuvCon | F(2,75) = 0.056 | .814 | .001 |
|  | Sex X NeoCon | F(1,75) = 0.175 | .840 | .005 |
|  | JuvCon X NeoCon | F(2,75) = 0.249 | .780 | .007 |
|  | Sex X JuvCon X NeoCon | F(2,75) = 0.426 | .654 | .011 |
| Crfr1 - Number |  |  |  |  |
|  | Hemisphere | F(1,74) = 0.527 | .470 | .007 |
|  | Hemisphere X Sex | F(1,74) = 1.799 | .184 | .024 |
|  | Hemi X Juv. Condition | F(1,74) = 3.829 | .054 | .049 |
|  | Hemi X Neo. Condition | F(2,74) = 1.967 | .147 | .050 |
|  | Hemi X Sex X JuvCon | F(1,74) = 0.153 | .697 | .002 |
|  | Hemi X Sex X NeoCon | F(2,74) = 0.786 | .460 | .021 |
|  | Hemi X JuvCon X NeoCon | F(2,74) = 1.109 | .335 | .029 |
|  | Hemi X sex X JuvCon X NeoCon | F(2,74) = 0.066 | .937 | .002 |
|  | Sex | F(1,74) = 0.031 | .861 | .000 |
|  | JuvCon | F(1,74) = 2.203 | .142 | .029 |
|  | NeoCon | F(2,74) = 0.160 | .853 | .004 |
|  | JuvCon X NeoCon | F(2,74) = 0.625 | .538 | .017 |
|  | Sex X JuvCon | F(1,74) = 0.133 | .717 | .002 |
|  | Sex X NeoCon | F(2,74) = 0.068 | .935 | .002 |
|  | Sex X JuvCon X NeoCon | F(2,74) = 0.075 | .928 | .002 |
| Crfr1 –<br>% cells |  |  |  |  |
| | Hemisphere | $F(1,75) = 2.20$ | .142 | .028 |
| | Hemisphere X Sex | $F(1,75) = 2.532$ | .116 | .033 |
| | Hemi X Juv. Condition | $F(1,75) = 1.023$ | .315 | .013 |
| | Hemi X Neo. Condition | $F(2,75) = 1.359$ | .263 | .035 |
| | Hemi X Sex X JuvCon | $F(1,75) = 0.241$ | .625 | .003 |
| | Hemi X Sex X NeoCon | $F(2,75) = 1.671$ | .195 | .043 |
| | Hemi X JuvCon X<br>NeoCon | $F(2,75) = 0.989$ | .377 | .026 |
| | Hemi X sex X JuvCon X<br>NeoCon | $F(2,75) = 1.668$ | .196 | .043 |
| | Sex | $F(1,75) = 0.006$ | .936 | .000 |
| | JuvCon | $F(1,75) = 0.023$ | .879 | .000 |
| | NeoCon | $F(2,75) = 0.045$ | .956 | .001 |
| | JuvCon X NeoCon | $F(2,75) = 1.253$ | .292 | .032 |
| | Sex X JuvCon X NeoCon | $F(2,75) = 0.376$ | .688 | .010 |
| Crfr1 –<br>% pixels |  |  |  |  |
| | Hemisphere | $F(1,74) = 0.868$ | .355 | .012 |
| | Hemisphere X Sex | $F(1,74) = 2.445$ | .122 | .032 |
| | Hemi X Juv. Condition | $F(1,74) = 0.025$ | .875 | .000 |
| | Hemi X Neo. Condition | $F(2,74) = 1.712$ | .188 | .054 |
| | Hemi X Sex X JuvCon | $F(1,74) = 0.201$ | .656 | .003 |
| | Hemi X Sex X NeoCon | $F(2,74) = 0.716$ | .492 | .019 |
| | Hemi X JuvCon X<br>NeoCon | $F(2,74) = 0.437$ | .648 | .012 |
| | Hemi X sex X JuvCon X<br>NeoCon | $F(2,74) = 2.074$ | .133 | .053 |
| | Sex | $F(1,74) = 0.439$ | .510 | .006 |
| | JuvCon | $F(1,74) = 0.405$ | .526 | .005 |
| | NeoCon | $F(2,74) = 0.156$ | .856 | .004 |
| | JuvCon X NeoCon | $F(2,74) = 0.102$ | .903 | .003 |
| | Sex X JuvCon | $F(1,74) = 2.987$ | .088 | .039 |
| | Sex X NeoCon | $F(2,74) = 0.001$ | .999 | .000 |
| | Sex X JuvCon X NeoCon | $F(2,74) = 0.535$ | .588 | .014 |
| Crfr2 -<br>Number |  |  |  |  |
| | Hemisphere | $F(1,71) = 5.325$ | **.024** | .070 |
| | Hemisphere X Sex | $F(1,71) = 1.183$ | .280 | .016 |
| | Hemi X Juv. Condition | $F(1,71) = 4.361$ | **.040** | .058 |
| | Hemi X Neo. Condition | $F(2,71) = 0.234$ | .792 | .007 |
| | Hemi X Sex X JuvCon | $F(1,71) = 1.290$ | .260 | .018 |
| | Hemi X Sex X NeoCon | $F(2,71) = 0.199$ | .820 | .006 |
| | Hemi X JuvCon X NeoCon | $F(2,71) = 0.336$ | .716 | .009 |
| | Hemi X sex X JuvCon X NeoCon | $F(2,71) = 1.804$ | .172 | .048 |
| | Sex | $F(1,71) = 1.594$ | .211 | .022 |
| | JuvCon | $F(1,71) = 0.061$ | .805 | .001 |
| | NeoCon | $F(2,71) = 0.662$ | .519 | .018 |
| | JuvCon X NeoCon | $F(2,71) = 0.249$ | .780 | .007 |
| | Sex X JuvCon | $F(1,71) = 0.665$ | .418 | .009 |
| | Sex X NeoCon | $F(2,71) = 0.727$ | .487 | .020 |
| | Sex X JuvCon X NeoCon | $F(2,72) = 0/006$ | .994 | .012 |
| Crfr2 - %Cells |  |  |  |  |
| | Hemisphere | $F(1,73) = 6.587$ | **.012** | .083 |
| | Hemisphere X Sex | $F(1,73) = 0.750$ | .389 | .010 |
| | Hemi X Juv. Condition | $F(1,73) = 4.130$ | **.046** | .054 |
| | Hemi X Neo. Condition | $F(2,73) = 0.111$ | .895 | .003 |
| | Hemi X Sex X JuvCon | $F(1,73) = 2.292$ | .134 | .030 |
| | Hemi X Sex X NeoCon | $F(2,73) = 0.658$ | .521 | .018 |
| | Hemi X JuvCon X NeoCon | $F(2,73) = 0.689$ | .505 | .019 |
| | Hemi X sex X JuvCon X NeoCon | $F(2,73) = 4.644$ | **.013** | .113 |
| | Sex | $F(1,73) = 0.431$ | .428 | .009 |
| | JuvCon | $F(1,73) = 0.063$ | .802 | .001 |
| | NeoCon | $F(2,73) = 0.055$ | .947 | .002 |
| | JuvCon X NeoCon | $F(2,73) = 0.196$ | .822 | .005 |
| | Sex X JuvCon | $F(1,73) = 1.040$ | .311 | .014 |
| | Sex X NeoCon | $F(2,73) = 0.756$ | .473 | .020 |
| | Sex X JuvCon X NeoCon | $F(2,73) = 1.549$ | .219 | .041 |
| Crfr2 - %Pixels |  |  |  |  |
| | Hemisphere | $F(1,74) = 0.631$ | .430 | .008 |
| | Hemisphere X Sex | $F(1,74) = 0.176$ | .676 | .002 |
| | Hemi X Juv. Condition | $F(1,74) = 0.038$ | .845 | .001 |
| | Hemi X Neo. Condition | $F(2,74) = 1.093$ | .340 | .029 |
| | Hemi X Sex X JuvCon | $F(1,74) = 0.487$ | .616 | .013 |
| | Hemi X Sex X NeoCon | $F(2,74) = 0.230$ | .795 | .006 |
| | Hemi X JuvCon X NeoCon | $F(2,74) = 0.487$ | .616 | .013 |
| | Hemi X sex X JuvCon X NeoCon | $F(2,74) = 4.455$ | .015 | .107 |
| | Sex | $F(1,74) = 0.147$ | .703 | .002 |
| | JuvCon | $F(1,74) = 0.219$ | .641 | .003 |
| | NeoCon | $F(2,74) = 1.247$ | .293 | .033 |
|  | JuvCon X NeoCon | F(2,74) = 0.115 | .892 | .003 |
|  | Sex X JuvCon | F(1,74) = 0.913 | .343 | .012 |
|  | Sex X NeoCon | F(2,74) = 1.486 | .233 | .039 |
|  | Sex X JuvCon X NeoCon | F(2,74) = 0.068 | .934 | .137 |

Number of cells expressing *crhr2*. The 2 (Sex: M,F) X 3 (Neonatal condition: undisturbed, handled, early pain) X 2 (Juvenile treatment: Fear Conditioned or not) X 2 (hemisphere) mixed model GLM (*Figure 8A*) yielded a significant main effect of hemisphere F(1,71) = 5.325, p<.05 η^2^ =.070 as well as an interaction between hemisphere and juvenile treatment F (1,71) = 4.36, p<.05 η^2^ =.058, with a significant increase in *crfr2* expression on the right side only in control subjects (p<.01; η^2^ =.226) (Figure 8B). There were no other significant statistical differences (see Table 2)

**Figure 8:**
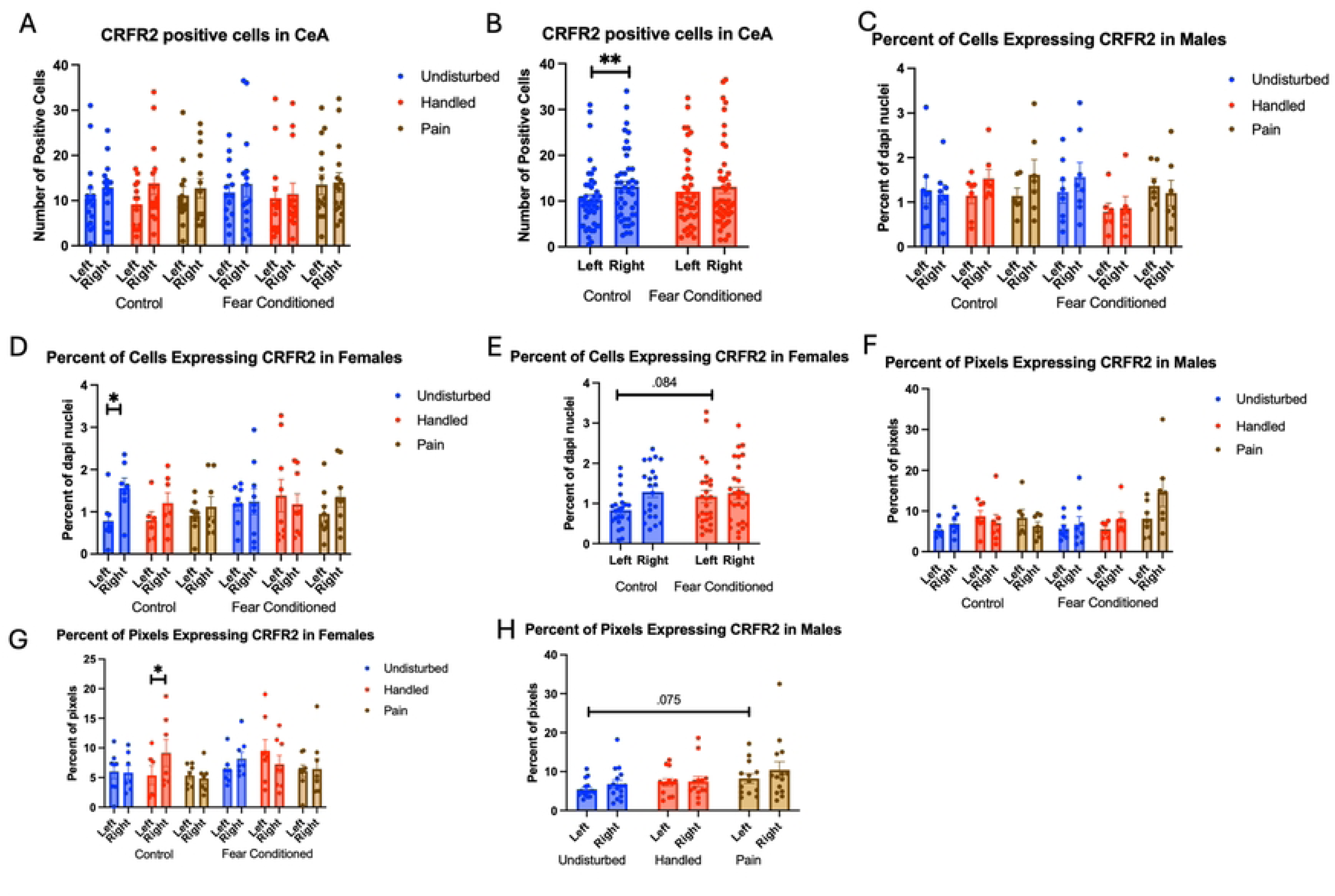
crhr2 expression in the CeA. PD 1-7 male and female rats were subjected to a painful experience, non-painful maternal separation and handling, or left undisturbed except for normal animal colony maintenance. Subjects were either fear conditioned or not on PD 24. Tissue was collected and subjected to RNAscope in situ hybridization in both hemispheres. Panel A: Number of cells expressing crhr2 from both hemispheres, collapsed across sex. Panel B: Number of cells expressing crhr2 collapsed across neonatal condition and sex, to highlight the hemisphere by juvenile condition statistical interaction. Panel C: Percentage of cells in overall ROI expressing crhr1 in males. Panel D: Percentage of cells in overall ROI expressing crhr2 in females. Panel E: Percentage of cells in overall ROI expressing crhr1 in females highlighting the hemisphere X juvenile condition interaction. Panel F: Percentage of pixels expressing positive signal after thresholding in males. Panel G: Percentage of pixels expressing positive signal after thresholding in females. Panel H: Percentage of pixels expressing positive signal after thresholding in males highlighting the trend towards an effect of sex. *= significant p<.05, **=p<.01.

The GLM examining the percentage of DAPI-stained cells expressing *crhr2* also revealed a significant main effect of hemisphere F(1,73) = 6.59, p<.05 η^2^ =.083, a significant hemisphere X juvenile treatment interaction F(1,73) = 4.13, p<.05 η^2^ =.054 and a significant 4-way interaction among hemisphere X sex X neonatal condition X juvenile treatment F (2, 73) = 4.64, p<.05 η^2^ =.113 (*Figures 8C – Males & 8D – Females)*. Follow up analysis found a trend towards a significant effect of juvenile treatment only on the left side of females (Figure 8E) F(1,38) = 3.155, p=.084 η^2^ =.077, suggesting that the statistical interaction is likely the result of minor hemispheric differences.

The GLM examining the percentage of pixels expressing *crhr2* found a significant 4-way interaction among hemisphere X juvenile condition X neonatal condition X sex F(2,74) = 4.455, p<.05 η^2^ =.107 (Figure 8F – Males and 8G – Females). Subsequent analyses demonstrate a significant hemisphere X juvenile condition X neonatal condition only in females F (2,38) = 4.978, p<.05 η^2^ =.208 and a trend towards a significant effect of Neonatal Condition only in left hemisphere of males F(2,36) = 2.628, p=.086, η^2^ =.127 (Figure 8H). This moderate-large effect on percent of pixels that express the target likely reflects changes in transcription within existing populations as well as recruitment of new populations of cells following early life pain.

### Periventricular Nucleus of the Hypothalamus

A total of 73-78 brains survived quality control and outlier analysis for the PVN for each measure. There was a minimum of 5 and a maximum of 8 subjects per group.

Number of nuclei stained with DAPI (Figure 9A). These data were again subjected to a 2 (Sex: M,F) X 3 (Neonatal condition undisturbed, handled, early pain) X 2 (Juvenile treatment: Fear Conditioned or not) X 2 (hemisphere) mixed model GLM for the number of stained nuclei. We again found an unexpected significant effect of hemisphere F(1,66) = 68.01, p<.001 η^2^ =.507 with the left side expressing more cells (Figure 9B).

**Figure 9:**
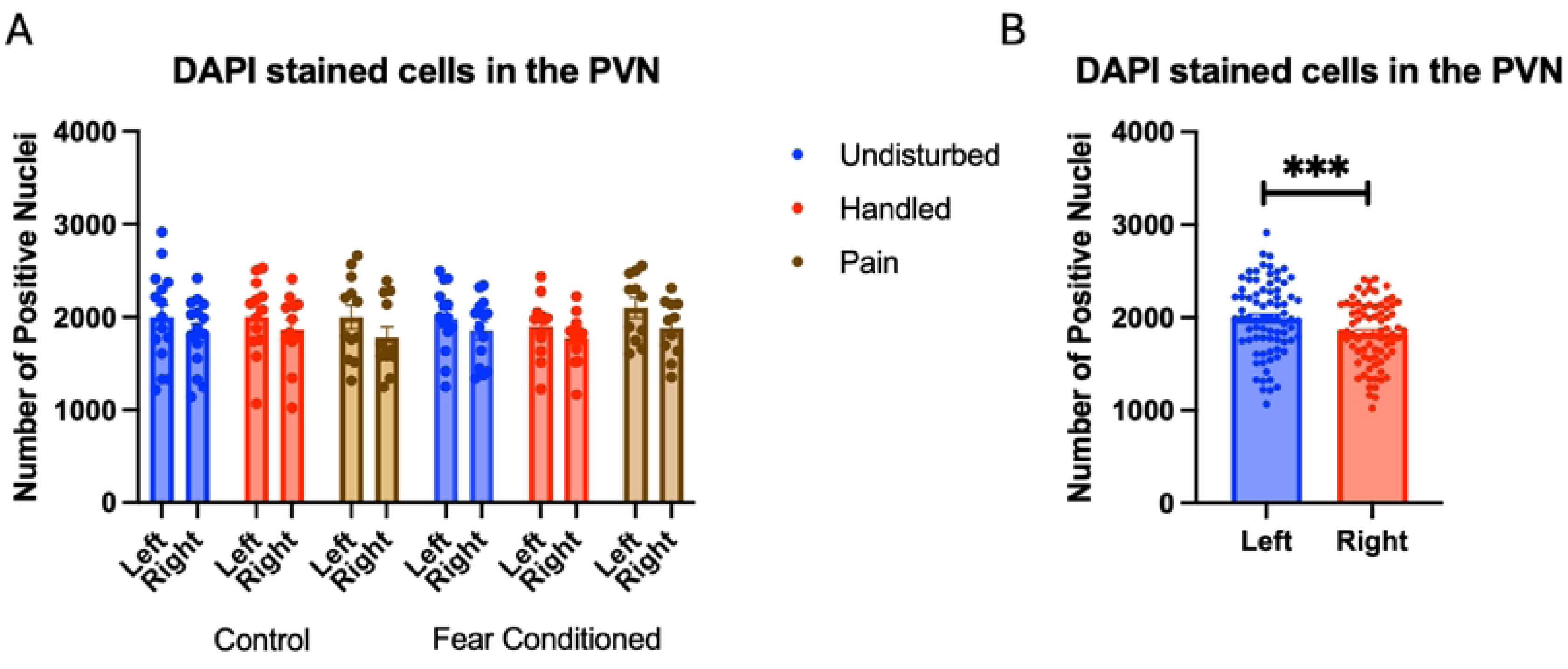
DAPI stained nuclei in the PVN. PD 1-7 male and female rats were subjected to a painful neonatal experience, non-painful removal from dam and handling, or left undisturbed except for normal animal colony procedures. Subjects were either fear conditioned or not on PD 24. Tissue was collected and subjected to RNAscope in situ hybridization in both hemisphere as shown in Panel A. There were significantly more DAPI-stained in the left hemisphere (Panel B). These differences were attributable to incidental differences in ROI size. *** = statistically significant p<.001.

Number of Cells expressing *crhr1*: As in the amygdala, these data were first subjected to a 2 (Sex: M,F) X 3 (Neonatal condition: undisturbed, handled, early pain) X 2 (Juvenile treatment: Fear Conditioned or not) X 2 (hemisphere) mixed model GLM for the number of cells expressing *crhr1* (Figure 10A - Males and 10B - Females). This analysis yielded a significant main effect of hemisphere F(1, 66) = 17.60, p<.001 η^2^ =.211 with the left hemisphere having more expression (Figure 10C), and a significant 4-way interaction among hemisphere, sex, neonatal, and juvenile condition F (2,66) = 5.54, p<.01 η^2^ =.144. When separated by hemisphere, there were no significant differences, suggesting that different patterns of hemispheric differences in males and females largely explain the 4 way interaction and these differences can largely be accounted for by differences in DAPI expression. There were no other significant effects or trends (see Table 3 for all results).

**Figure 10:**
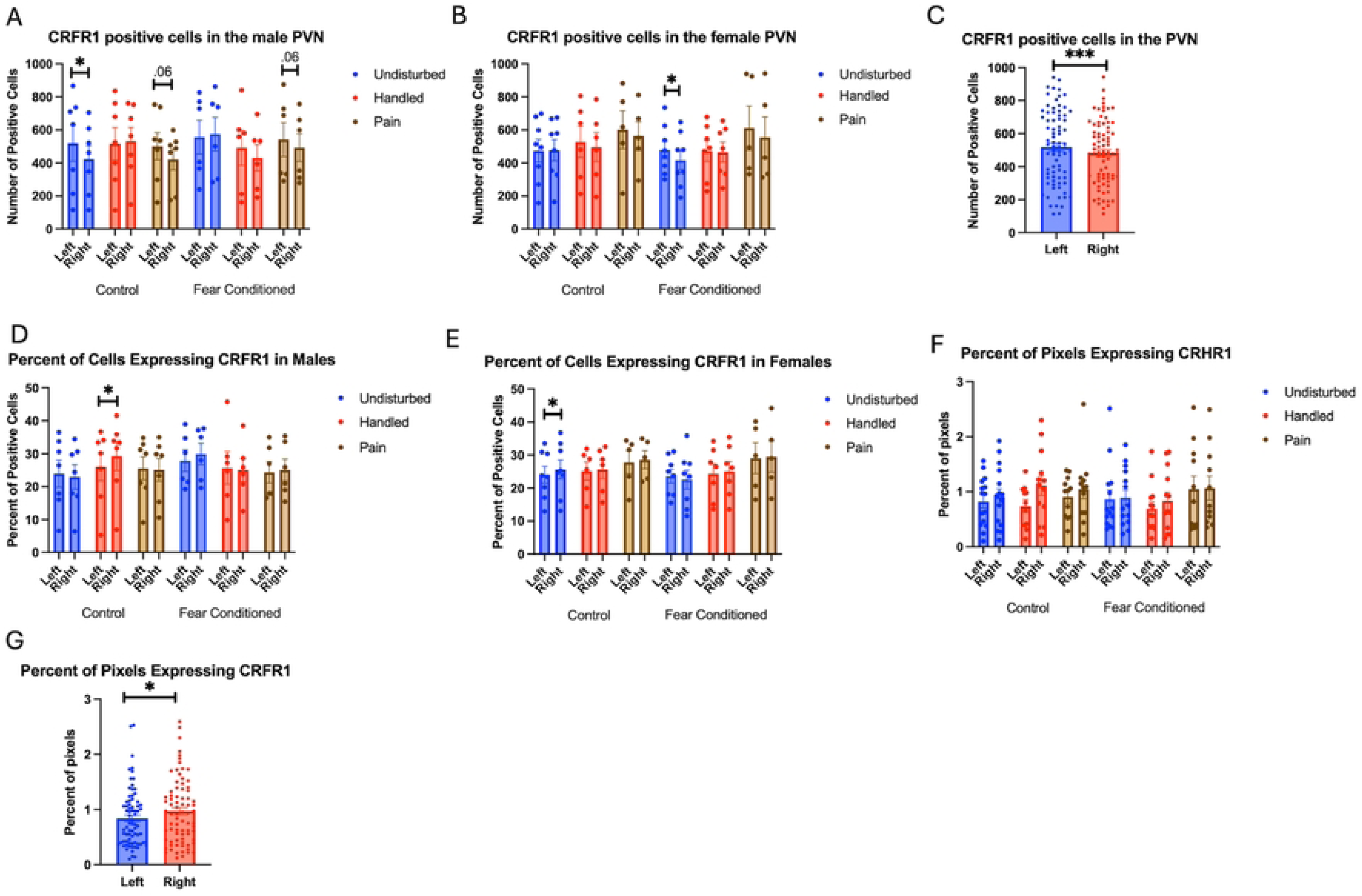
crhr1 expression in the PVN. PD 1-7 male and female rats were subjected to a painful experience, non-painful maternal separation and handling, or left undisturbed except for normal animal colony maintenance. Subjects were either fear conditioned or not on PD 24. Tissue was collected and subjected to RNAscope in situ hybridization in both hemispheres. Panel A: Number of cells expressing crhr1 from both hemispheres, in males. Panel B: Number of cells expressing crhr1 from both hemispheres, in females. Panel C: Number of cells expressing crhr1 demonstrating the effects of hemisphere. Panel D: Percentage of cells expressing crhr1 in males. Panel E: Percentage of cells expressing crhr1 in females. Panel F: Percentage of pixels expressing crhr1 after thresholding. Panel G: Percentage of pixels expressing crhr1 highlighting the hemisphere effect. * = statistically significant p<.05 *** = p<.001.

**Table 3:**
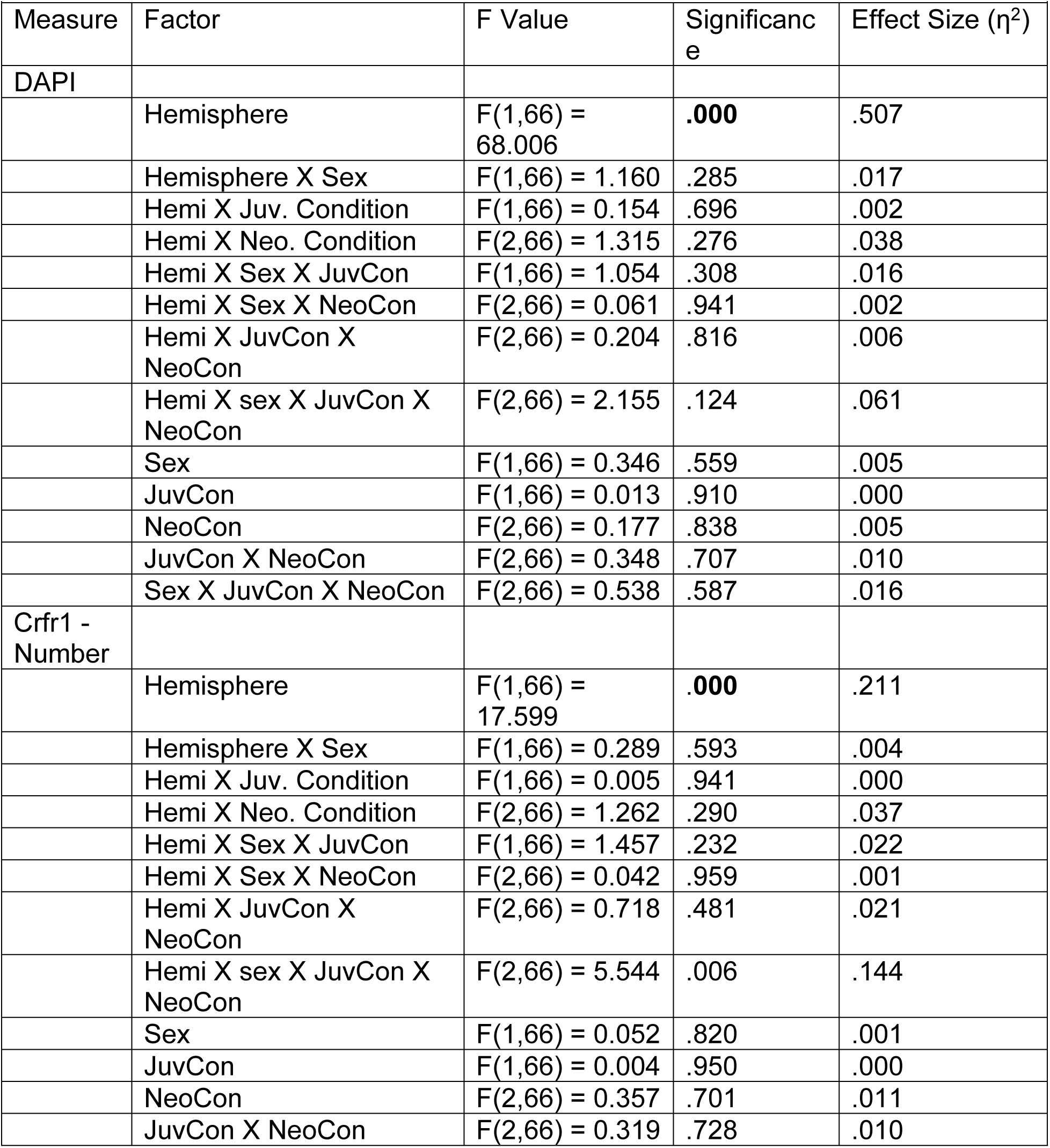
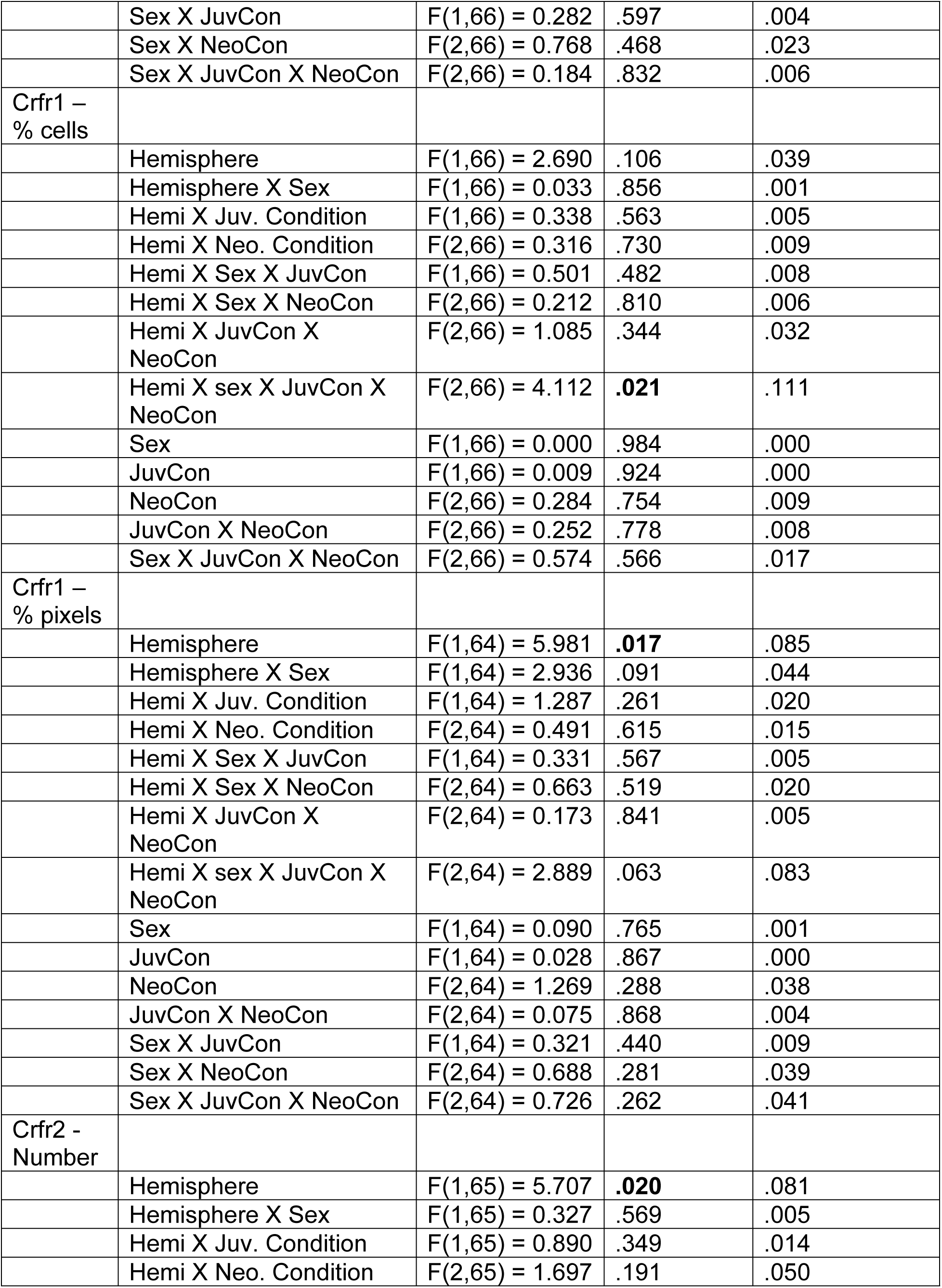
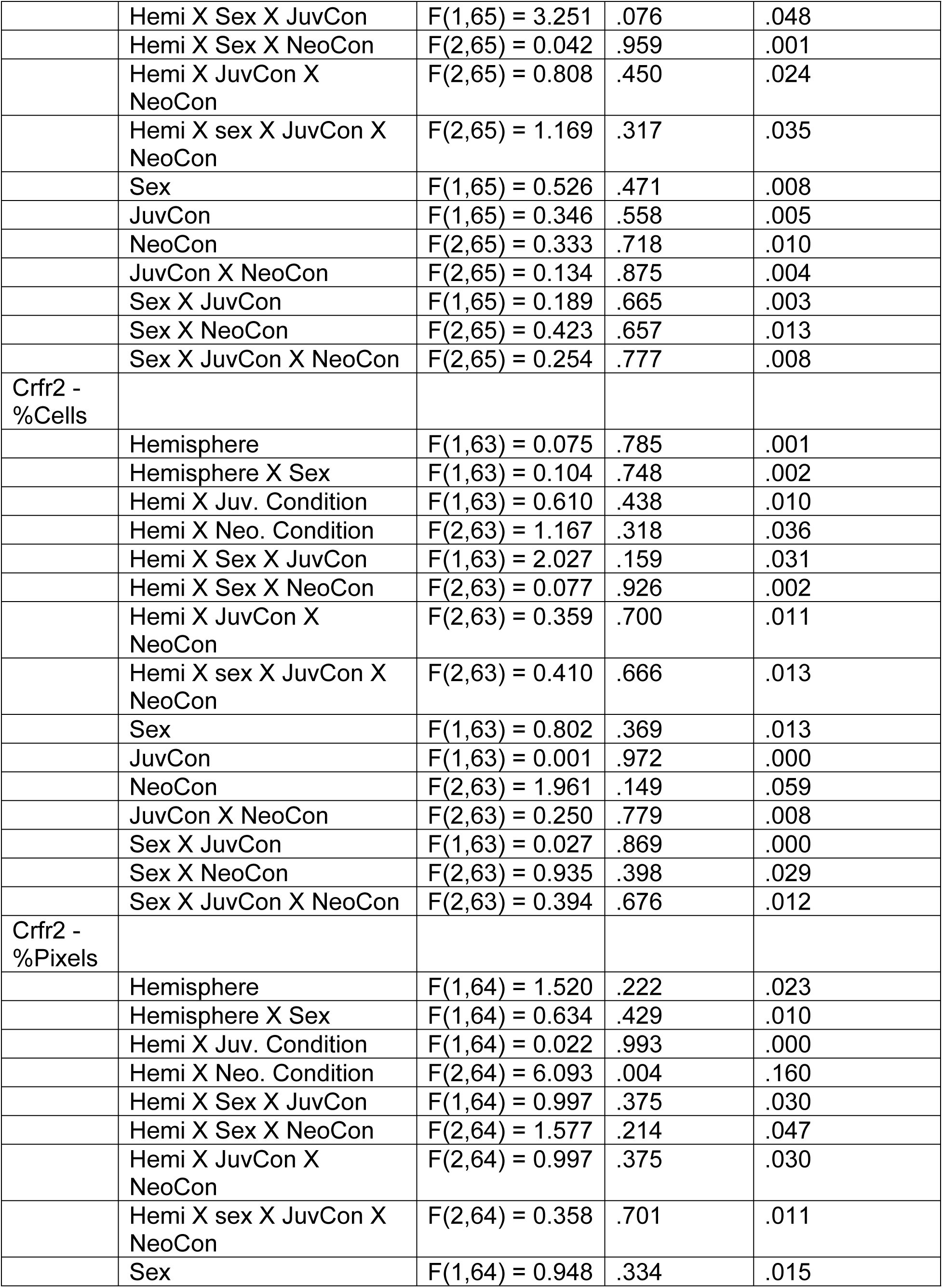
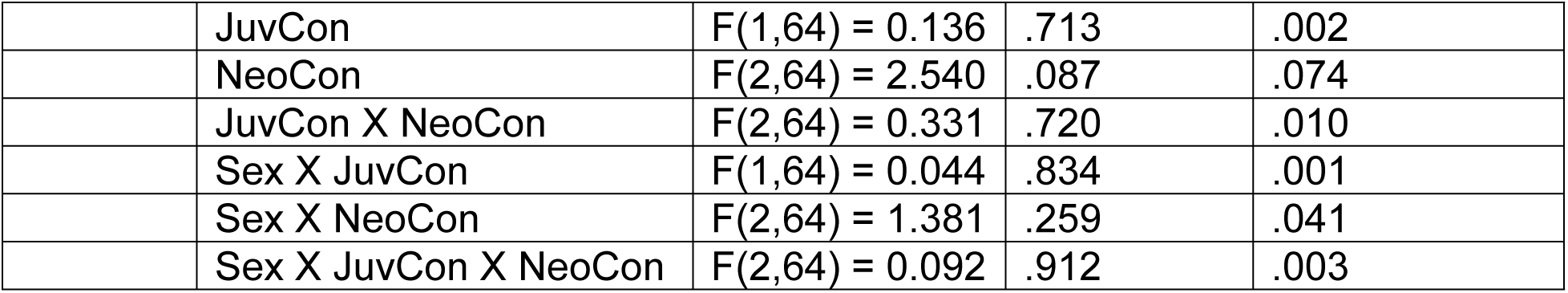
PVN.

When examining the percent of DAPI stained cells that express *crhr1,* we find a significant 4-way interaction among hemisphere, sex, juvenile condition, and neonatal condition F(2,66) = 4.112, p<.05 η^2^ =.111 (Figure 10D – Males 10E – Females). This appears to be attributable to minor hemisphere differences between the sexes, as follow-up analyses separated by the other variables find no other significant differences as a function of sex, juvenile condition, or neonatal condition (all analyses found in Table 3).

An analysis examining percent of pixels expressing *crhr1* demonstrated a significant main effect of hemisphere F(1,64) = 5.981 p<.05 η^2^ =.085 (Figure 10F), attributable to increases in expression in the right hemisphere (Figure 10G). There were no other significant effects (all analyses found in Table 3).

Number of Cells expressing *crhr2* (Figure 11A): The 4-way analysis found a significant effect of hemisphere F(1,65) = 5.71, p<.05, η^2^ =.081 with greater expression on the left side (Figure 11B), with no other significant effects (see Table 3). Examining the percent of DAPI-stained cells expressing *crhr2 (*Figure 11C) demonstrated no significant effects or interactions.

**Figure 11:**
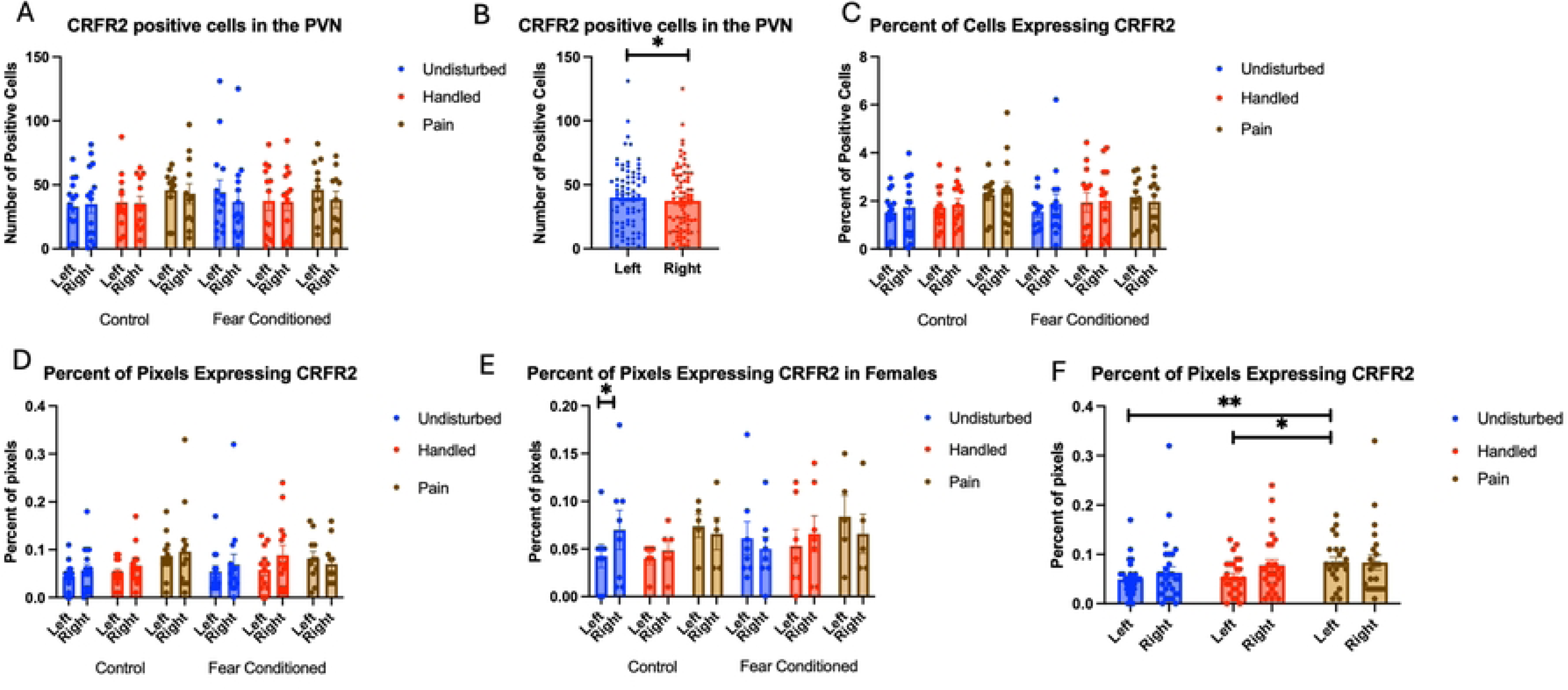
crhr2 expression in the PVN. PD 1-7 male and female rats were subjected to a painful experience, non-painful maternal separation and handling, or left undisturbed except for normal animal colony maintenance. Subjects were either fear conditioned or not on PD 24. Tissue was collected and subjected to RNAscope in situ hybridization in both hemispheres. Panel A: Number of cells expressing crhr2 from both hemispheres, collapsed across sex. Panel B: Number of cells expressing crhr2 to highlight the effect of hemisphere. Panel C: Percentage of cells in overall ROI expressing crhr2. Panel D: Percentage of pixels expressing positive signal after thresholding. Panel E: Percentage of pixels expressing positive signal after thresholding in females. Panel F: Percentage of pixels expressing positive signal after thresholding highlighting the effects of neonatal pain. *= significant p<.05, **=p<.01.

Examining the percent of pixels expressing *crhr2* (Figure 11D) finds a significant interaction between hemisphere and neonatal condition F(2,64) = 6.093, p<.01 η^2^ =.160, and a 3-way interaction among hemisphere, sex and juvenile condition F(1,64) = 4.017, p<.05, η^2^ =.059. Further analyses demonstrate an effect of hemisphere only in female, undisturbed, control subjects (Figure 11E). An ANOVA on expression in the left hemisphere finds a significant effect of neonatal condition F(2,64) = 5.385, p<.01 η^2^ =.144 (a large effect size) with the neonatal pain group differing significantly from the other two groups (Figure 11F). There were no significant differences on the right, despite a similar pattern. The overall pattern suggests that neonatal pain increases expression of *crhr2* receptors in the PVN without altering the number of cells that express them.

### Ventromedial Hypothalamus

A total of 75-78 brains survived quality control and outlier analysis for the VMH for each measure. There was a minimum of 5 and a maximum of 8 subjects per group.

Number of nuclei stained with DAPI (Figure 12A). These data were again subjected to a 2 (Sex: M,F) X 3 (Neonatal treatment: undisturbed, handled, early pain) X 2 (Juvenile treatment: Fear Conditioned or not) X 2 (hemisphere) mixed model GLM for the number of stained nuclei. We again found an effect of hemisphere F(1,66) = 23.242, p<.001 η^2^ =.260 with the left side expressing more cells (Figure 12B). All analyses can be found in Table 4.

**Figure 12:**
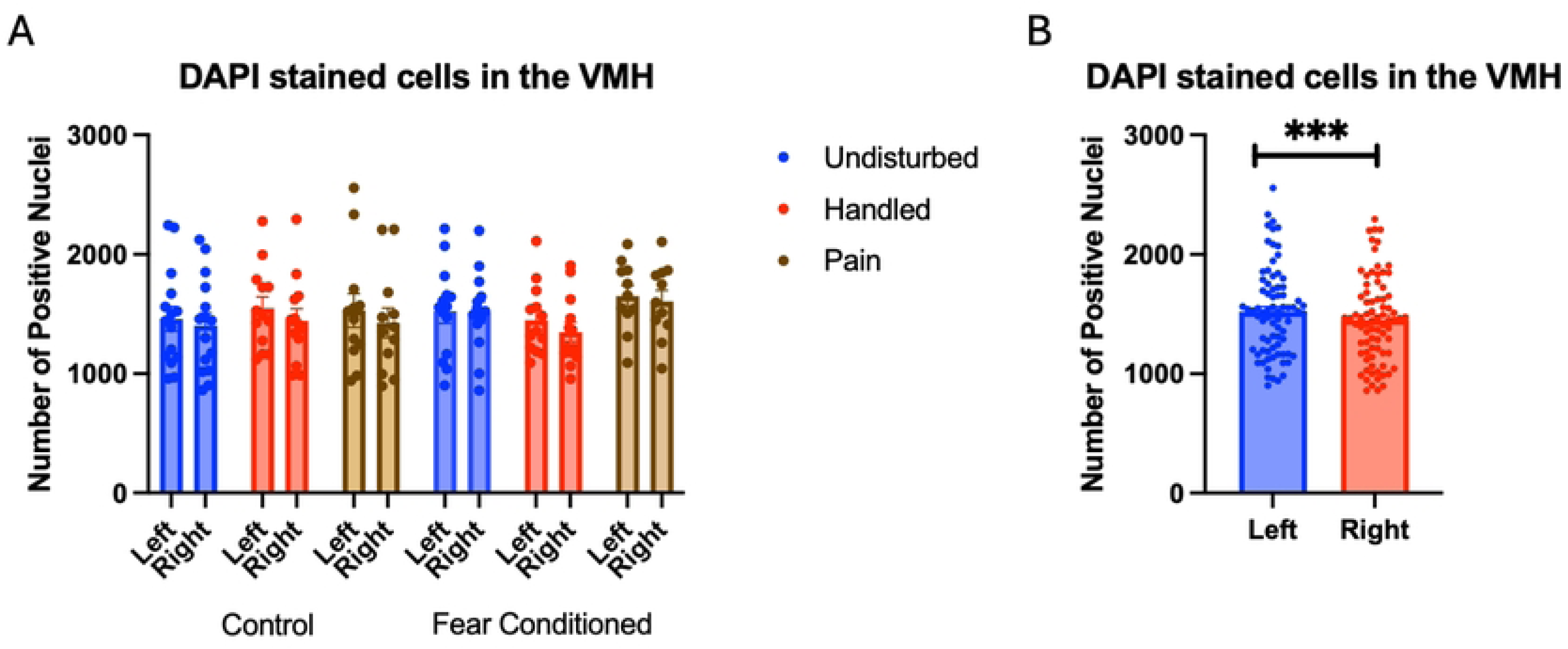
DAPI stained nuclei in the VMH. PD 1-7 male and female rats were subjected to a painful neonatal experience, non-painful maternal separation and handling, or left undisturbed except for normal animal colony procedures. Subjects were either fear conditioned or not on PD 24. Tissue was collected and subjected to RNAscope in situ hybridization in both hemisphere as shown in Panel A. There were significantly more DAPI-stained in the left hemisphere (Panel B). These differences were attributable to incidental differences in ROI size. *** = statistically significant p<.001.

**Table 4:**
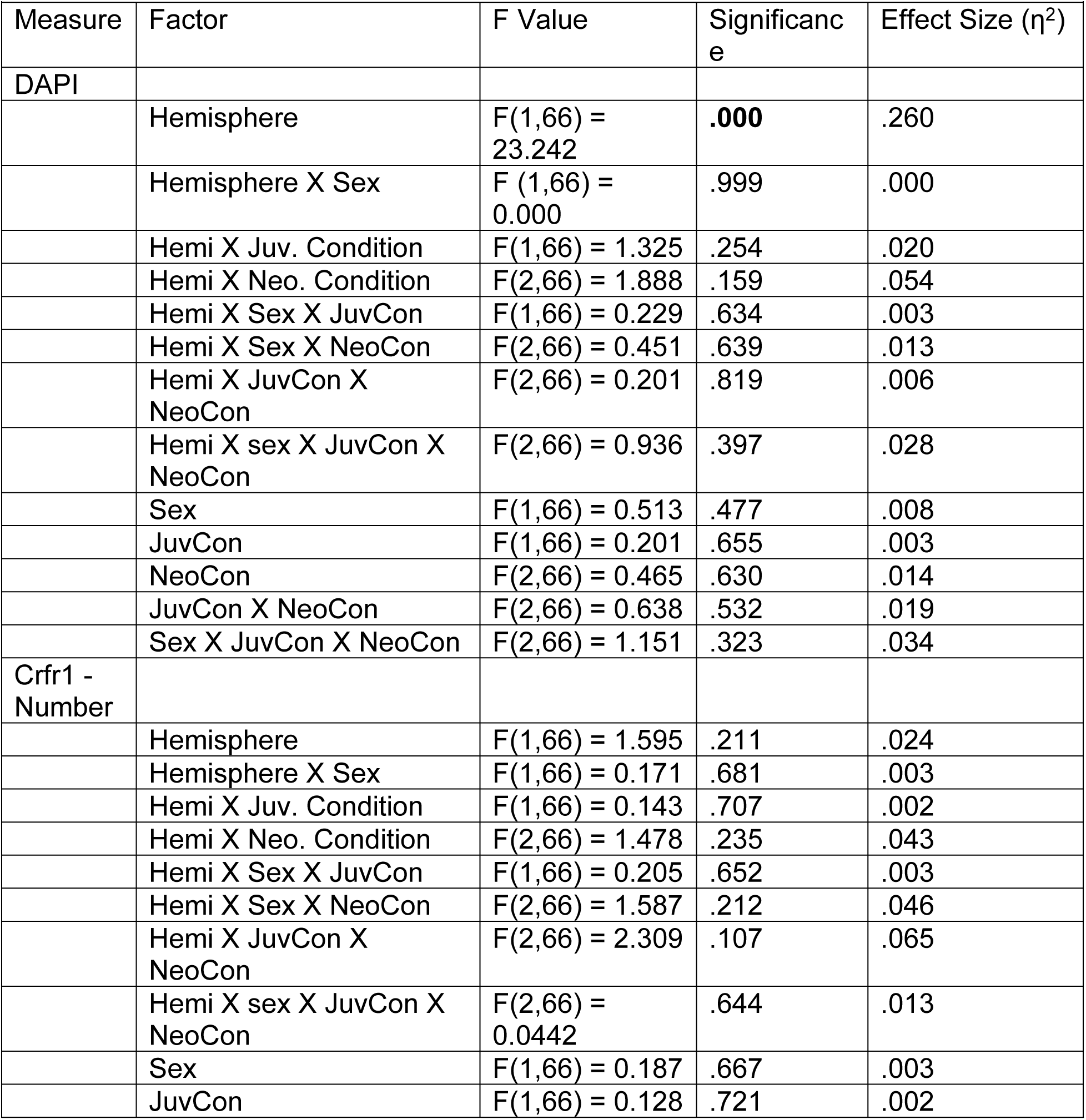
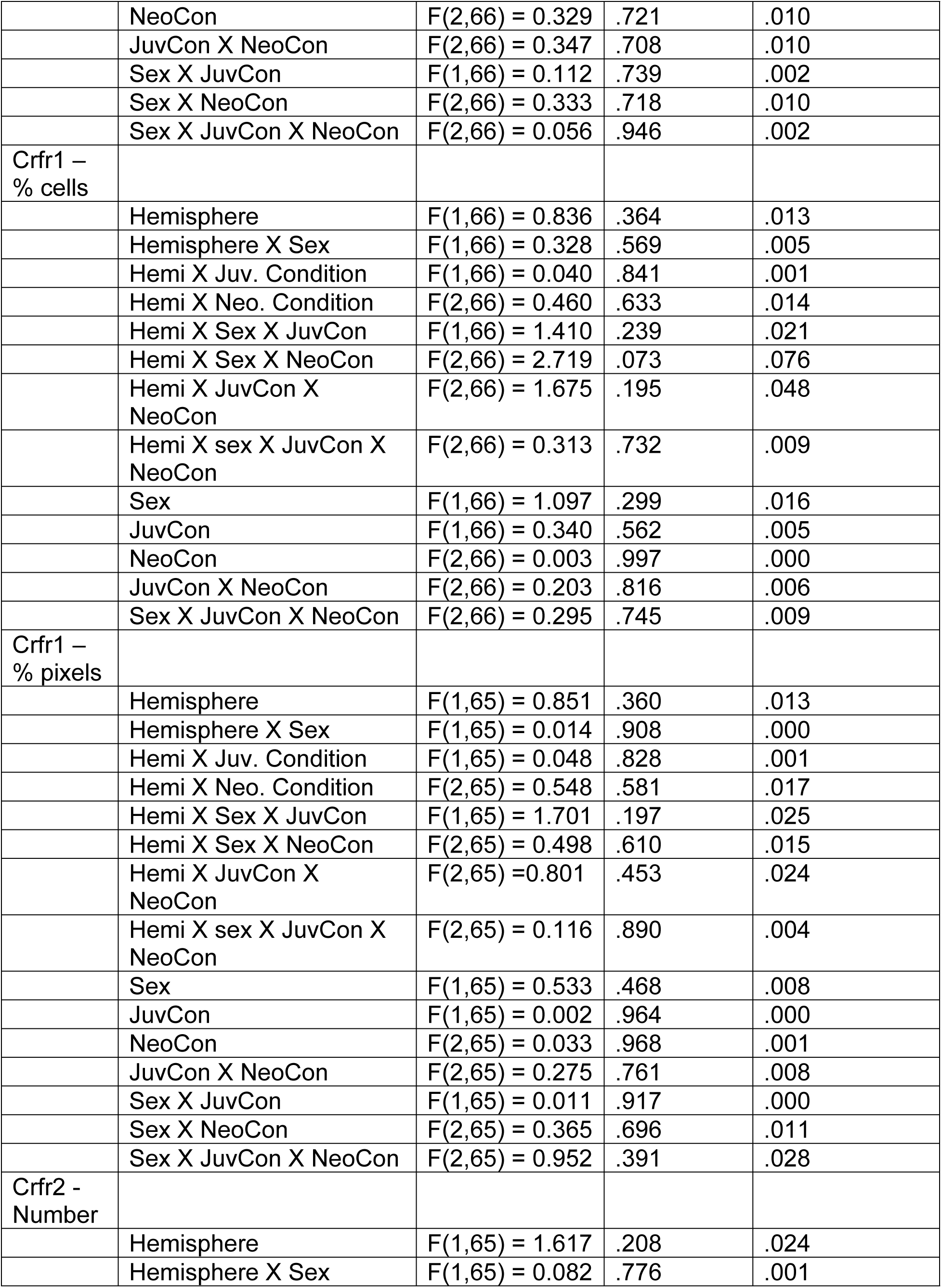
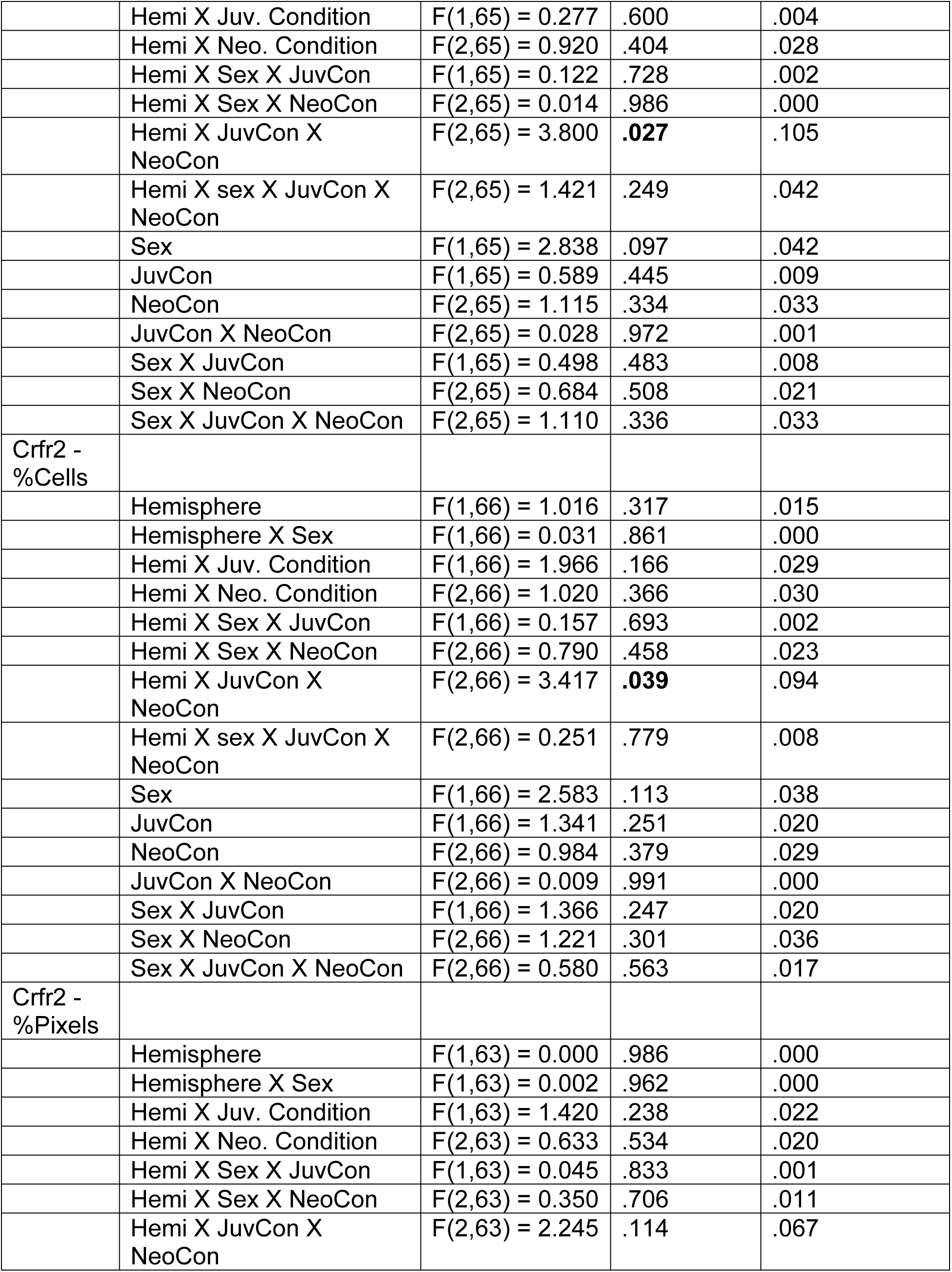
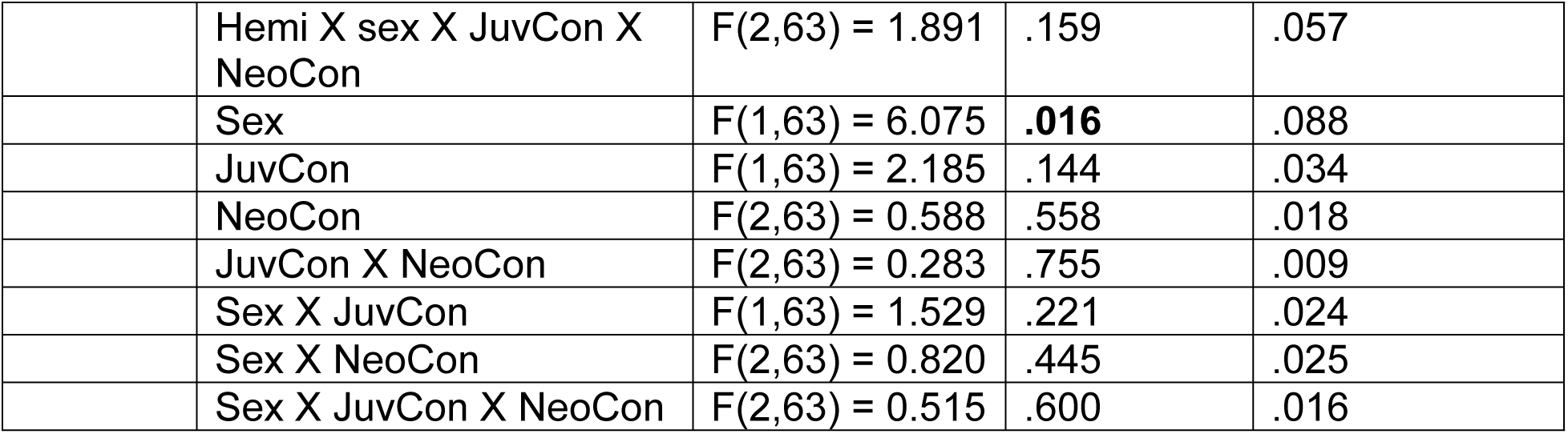
VMH.

| Table 4: VMH |  |  |  |  |
| --- | --- | --- | --- | --- |
| Measure | Factor | F Value | Significance | Effect Size ($\eta^2$) |
| DAPI |  |  |  |  |
|  | Hemisphere | F(1,66) = 23.242 | **.000** | .260 |
|  | Hemisphere X Sex | F (1,66) = 0.000 | .999 | .000 |
|  | Hemi X Juv. Condition | F(1,66) = 1.325 | .254 | .020 |
|  | Hemi X Neo. Condition | F(2,66) = 1.888 | .159 | .054 |
|  | Hemi X Sex X JuvCon | F(1,66) = 0.229 | .634 | .003 |
|  | Hemi X Sex X NeoCon | F(2,66) = 0.451 | .639 | .013 |
|  | Hemi X JuvCon X NeoCon | F(2,66) = 0.201 | .819 | .006 |
|  | Hemi X sex X JuvCon X NeoCon | F(2,66) = 0.936 | .397 | .028 |
|  | Sex | F(1,66) = 0.513 | .477 | .008 |
|  | JuvCon | F(1,66) = 0.201 | .655 | .003 |
|  | NeoCon | F(2,66) = 0.465 | .630 | .014 |
|  | JuvCon X NeoCon | F(2,66) = 0.638 | .532 | .019 |
|  | Sex X JuvCon X NeoCon | F(2,66) = 1.151 | .323 | .034 |
| Crfr1 - Number |  |  |  |  |
|  | Hemisphere | F(1,66) = 1.595 | .211 | .024 |
|  | Hemisphere X Sex | F(1,66) = 0.171 | .681 | .003 |
|  | Hemi X Juv. Condition | F(1,66) = 0.143 | .707 | .002 |
|  | Hemi X Neo. Condition | F(2,66) = 1.478 | .235 | .043 |
|  | Hemi X Sex X JuvCon | F(1,66) = 0.205 | .652 | .003 |
|  | Hemi X Sex X NeoCon | F(2,66) = 1.587 | .212 | .046 |
|  | Hemi X JuvCon X NeoCon | F(2,66) = 2.309 | .107 | .065 |
|  | Hemi X sex X JuvCon X NeoCon | F(2,66) = 0.0442 | .644 | .013 |
|  | Sex | F(1,66) = 0.187 | .667 | .003 |
|  | JuvCon | F(1,66) = 0.128 | .721 | .002 |
| | NeoCon | $F(2,66) = 0.329$ | .721 | .010 |
| | JuvCon X NeoCon | $F(2,66) = 0.347$ | .708 | .010 |
| | Sex X JuvCon | $F(1,66) = 0.112$ | .739 | .002 |
| | Sex X NeoCon | $F(2,66) = 0.333$ | .718 | .010 |
| | Sex X JuvCon X NeoCon | $F(2,66) = 0.056$ | .946 | .002 |
| Crfr1 –<br>% cells |  |  |  |  |
| | Hemisphere | $F(1,66) = 0.836$ | .364 | .013 |
| | Hemisphere X Sex | $F(1,66) = 0.328$ | .569 | .005 |
| | Hemi X Juv. Condition | $F(1,66) = 0.040$ | .841 | .001 |
| | Hemi X Neo. Condition | $F(2,66) = 0.460$ | .633 | .014 |
| | Hemi X Sex X JuvCon | $F(1,66) = 1.410$ | .239 | .021 |
| | Hemi X Sex X NeoCon | $F(2,66) = 2.719$ | .073 | .076 |
| | Hemi X JuvCon X<br>NeoCon | $F(2,66) = 1.675$ | .195 | .048 |
| | Hemi X sex X JuvCon X<br>NeoCon | $F(2,66) = 0.313$ | .732 | .009 |
| | Sex | $F(1,66) = 1.097$ | .299 | .016 |
| | JuvCon | $F(1,66) = 0.340$ | .562 | .005 |
| | NeoCon | $F(2,66) = 0.003$ | .997 | .000 |
| | JuvCon X NeoCon | $F(2,66) = 0.203$ | .816 | .006 |
| | Sex X JuvCon X NeoCon | $F(2,66) = 0.295$ | .745 | .009 |
| Crfr1 –<br>% pixels |  |  |  |  |
| | Hemisphere | $F(1,65) = 0.851$ | .360 | .013 |
| | Hemisphere X Sex | $F(1,65) = 0.014$ | .908 | .000 |
| | Hemi X Juv. Condition | $F(1,65) = 0.048$ | .828 | .001 |
| | Hemi X Neo. Condition | $F(2,65) = 0.548$ | .581 | .017 |
| | Hemi X Sex X JuvCon | $F(1,65) = 1.701$ | .197 | .025 |
| | Hemi X Sex X NeoCon | $F(2,65) = 0.498$ | .610 | .015 |
| | Hemi X JuvCon X<br>NeoCon | $F(2,65) = 0.801$ | .453 | .024 |
| | Hemi X sex X JuvCon X<br>NeoCon | $F(2,65) = 0.116$ | .890 | .004 |
| | Sex | $F(1,65) = 0.533$ | .468 | .008 |
| | JuvCon | $F(1,65) = 0.002$ | .964 | .000 |
| | NeoCon | $F(2,65) = 0.033$ | .968 | .001 |
| | JuvCon X NeoCon | $F(2,65) = 0.275$ | .761 | .008 |
| | Sex X JuvCon | $F(1,65) = 0.011$ | .917 | .000 |
| | Sex X NeoCon | $F(2,65) = 0.365$ | .696 | .011 |
| | Sex X JuvCon X NeoCon | $F(2,65) = 0.952$ | .391 | .028 |
| Crfr2 -<br>Number |  |  |  |  |
| | Hemisphere | $F(1,65) = 1.617$ | .208 | .024 |
| | Hemisphere X Sex | $F(1,65) = 0.082$ | .776 | .001 |
| | Hemi X Juv. Condition | $F(1,65) = 0.277$ | .600 | .004 |
| | Hemi X Neo. Condition | $F(2,65) = 0.920$ | .404 | .028 |
| | Hemi X Sex X JuvCon | $F(1,65) = 0.122$ | .728 | .002 |
| | Hemi X Sex X NeoCon | $F(2,65) = 0.014$ | .986 | .000 |
| | Hemi X JuvCon X NeoCon | $F(2,65) = 3.800$ | **.027** | .105 |
| | Hemi X sex X JuvCon X NeoCon | $F(2,65) = 1.421$ | .249 | .042 |
| | Sex | $F(1,65) = 2.838$ | .097 | .042 |
| | JuvCon | $F(1,65) = 0.589$ | .445 | .009 |
| | NeoCon | $F(2,65) = 1.115$ | .334 | .033 |
| | JuvCon X NeoCon | $F(2,65) = 0.028$ | .972 | .001 |
| | Sex X JuvCon | $F(1,65) = 0.498$ | .483 | .008 |
| | Sex X NeoCon | $F(2,65) = 0.684$ | .508 | .021 |
| | Sex X JuvCon X NeoCon | $F(2,65) = 1.110$ | .336 | .033 |
| Crfr2 - %Cells |  |  |  |  |
| | Hemisphere | $F(1,66) = 1.016$ | .317 | .015 |
| | Hemisphere X Sex | $F(1,66) = 0.031$ | .861 | .000 |
| | Hemi X Juv. Condition | $F(1,66) = 1.966$ | .166 | .029 |
| | Hemi X Neo. Condition | $F(2,66) = 1.020$ | .366 | .030 |
| | Hemi X Sex X JuvCon | $F(1,66) = 0.157$ | .693 | .002 |
| | Hemi X Sex X NeoCon | $F(2,66) = 0.790$ | .458 | .023 |
| | Hemi X JuvCon X NeoCon | $F(2,66) = 3.417$ | **.039** | .094 |
| | Hemi X sex X JuvCon X NeoCon | $F(2,66) = 0.251$ | .779 | .008 |
| | Sex | $F(1,66) = 2.583$ | .113 | .038 |
| | JuvCon | $F(1,66) = 1.341$ | .251 | .020 |
| | NeoCon | $F(2,66) = 0.984$ | .379 | .029 |
| | JuvCon X NeoCon | $F(2,66) = 0.009$ | .991 | .000 |
| | Sex X JuvCon | $F(1,66) = 1.366$ | .247 | .020 |
| | Sex X NeoCon | $F(2,66) = 1.221$ | .301 | .036 |
| | Sex X JuvCon X NeoCon | $F(2,66) = 0.580$ | .563 | .017 |
| Crfr2 - %Pixels |  |  |  |  |
| | Hemisphere | $F(1,63) = 0.000$ | .986 | .000 |
| | Hemisphere X Sex | $F(1,63) = 0.002$ | .962 | .000 |
| | Hemi X Juv. Condition | $F(1,63) = 1.420$ | .238 | .022 |
| | Hemi X Neo. Condition | $F(2,63) = 0.633$ | .534 | .020 |
| | Hemi X Sex X JuvCon | $F(1,63) = 0.045$ | .833 | .001 |
| | Hemi X Sex X NeoCon | $F(2,63) = 0.350$ | .706 | .011 |
| | Hemi X JuvCon X NeoCon | $F(2,63) = 2.245$ | .114 | .067 |
| | Hemi X sex X JuvCon X NeoCon | $F(2,63) = 1.891$ | .159 | .057 |
| | Sex | $F(1,63) = 6.075$ | **.016** | .088 |
| | JuvCon | $F(1,63) = 2.185$ | .144 | .034 |
| | NeoCon | $F(2,63) = 0.588$ | .558 | .018 |
| | JuvCon X NeoCon | $F(2,63) = 0.283$ | .755 | .009 |
| | Sex X JuvCon | $F(1,63) = 1.529$ | .221 | .024 |
| | Sex X NeoCon | $F(2,63) = 0.820$ | .445 | .025 |
| | Sex X JuvCon X NeoCon | $F(2,63) = 0.515$ | .600 | .016 |

Number of Cells expressing *crhr1*: 2 (Sex: M, F) X 3 (Neonatal condition: undisturbed, handled, early pain) X 2 (Juvenile treatment: Fear Conditioned or not) X 2 (hemisphere) mixed model GLMs for the number of cells (Figure13A), the percentage of cells (Figure 13B), or the percentage of pixels (Figure 13C) expressing *crhr1* yielded no significant effects or interactions. Detailed statistics can be found in Table 4.

**Figure 13:**
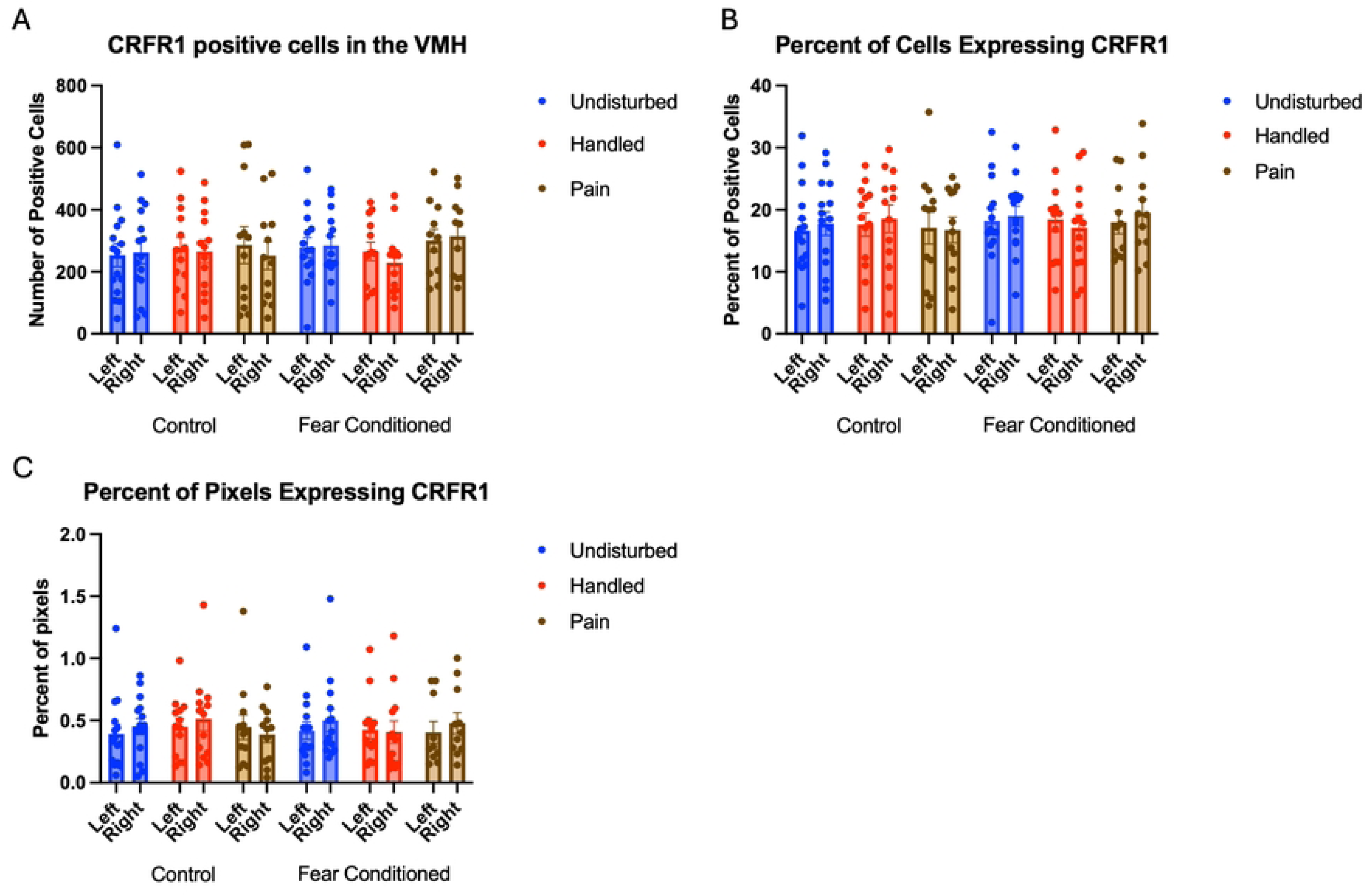
crhr1 expression in the VMH. PD 1-7 male and female rats were subjected to a painful experience, non-painful maternal separation and handling, or left undisturbed except for normal animal colony maintenance. Subjects were either fear conditioned or not on PD 24. Tissue was collected and subjected to RNAscope in situ hybridization in both hemispheres. Panel A: Number of cells expressing crhr1 from both hemispheres. Panel B: Percentage of DAPI-labeled cells expressing crhr1 from both hemispheres. Panel C: Percentage of pixels expressing crhr1 after thresholding. There were no significant differences.

For the number of cells expressing *CRFR2*, the GLM yielded a significant hemisphere X neonatal X juvenile treatment 3-way interaction F (2, 65) = 3.80, p<.05 η^2^ =.105 (Figure 14A). Further analyses show this is reflective of a minor hemispheric difference only in handled, fear conditioned subjects F(1,11) = 5.439, p<.05 η^2^ =.33. There were no other significant effects of neonatal condition or juvenile treatment.

**Figure 14:**
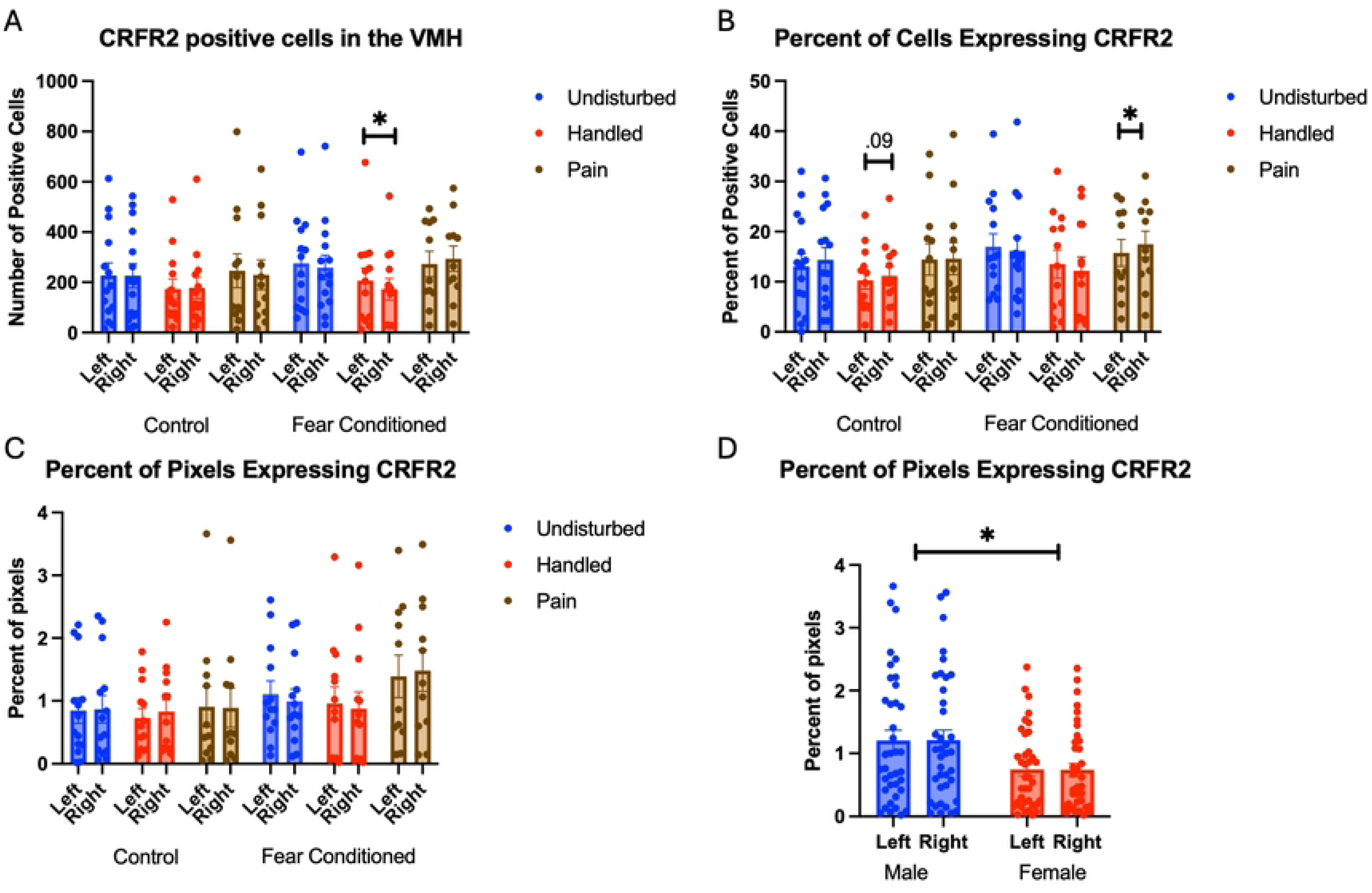
crhr2 expression in the VMH. PD 1-7 male and female rats were subjected to a painful experience, non-painful removal from dam and handling, or left undisturbed except for normal animal colony maintenance. Subjects were either fear conditioned or not on PD 24. Tissue was collected and subjected to RNAscope in situ hybridization in both hemispheres. Panel A: Number of cells expressing crhr2 from both hemispheres, collapsed across sex. Panel B: Percentage of cells in overall ROI expressing crhr2. Panel C: Percentage of pixels expressing positive signal after thresholding. Panel D: Percentage of pixels expressing positive signal highlighting the effect of sex. *= significant p<.05.

Analyses on the percent of DAPI-labeled cells (Figure 14B) expressing *crhr2* also yielded a significant hemisphere X neonatal X juvenile treatment 3-way interaction F(2.66) = 3.417, p<.05 η^2^ =.09 that was again reflective of minor hemispheric differences in neonatal handled, control subjects as well as neonatal pain fear conditioned subjects. The analysis on percent of pixels expressing *crhr2* (Figure 14C) yielding a significant main effect of sex F(1,63) = 8.762, p<.05 η^2^ =.09 with males demonstrating more expression than females (Figure 14D). See Table 4 for full statistics.

## Discussion

This paper examines the effects of both neonatal pain and juvenile stress on the expression of CRF receptor mRNA in critical regions of the amygdala and hypothalamus. Our prior work has demonstrated effects of our early life pain model on expression of *crh* mRNA in the CEA of only male (but not female) rats [46,47]. Thus, based on these findings and the literature, we anticipated that the CeA, BLA and PVN would show significant changes in receptor expression. The VMH was largely a region of convenience and serves as a control region showing the specificity of our effects in the critical regions. We were therefore surprised to find few effects of this model on CRH receptor expression in the amygdala and it suggests that the effects of early life pain are limited to *crh-*expressing cells.

In the current study, the early life pain experience had no meaningful effect on CRF receptor 1 expression within the amygdala and hypothalamus as assessed in the juvenile period. This is relatively surprising given prior data that a single inflammatory injection of carrageenan within a day of birth was found to significantly reduce CRFR1 expression in the BLA (among other changes) [61]. Importantly, we have previously shown that carrageenan injections do not produce the same effects as our repeated paw prick model [31,63], likely due to differences in duration and intensity of the pain, as well as the concomitant inflammation that accompanies carrageenan. This may account for some of the difference in outcomes, as might the fact that juveniles were assessed in the current study whereas Victoria et al. assessed adults.

We did observe increases in CRF receptor 2 expression following early life pain, especially in the PVN and less so in the CeA. CRF receptors 1 and 2 may play opposite, or at least differential, roles in emotional outcome, such as anxiety, fear, pain, and threat responding [67–70]. That the effect in the CeA was limited to males is consistent with our prior work, demonstrating effects on CRF-expressing cells in our model in the CeA only in male subjects [46,47], thus male-only effects on the CeA may be expected. Regarding the specificity of the effect to CRF receptor 2, there are known differences in CRF receptor function. CRF receptor 1 activation in the amygdala tends to escalate active forms of anxiety, as well as pain. In contrast, CRF receptor 2 appears to be involved in pain and stress resolution, or passive forms of anxiety and threat responding. Increases in *crhr2* is typically is associated with reductions in anxiety [69].

In our hands, this was an increase in overall expression, but not in increase in the number of cells showing expression, suggesting an upregulation in *crfr2* expression in the population of cells already expressing the receptor. Given that our prior work has demonstrated an anxiolytic effect of our model – decreasing conditioned freezing and increased time in open arms on the elevated plus maze [31,47], these changes in receptor expression appear to be consistent with the observed changes in behavior. Similar results have also been associated with chronic pain conditions [71]. Thus, when compared with our prior work and the literature, it appears that the NICU-like experience produces and lasting changes in CRF-expressing cells (e.g. Davis et al., 2021), but as well as changes in CRF-receptor 2 expression. Of course, the current data are not informative regarding any acute effects of the neonatal trauma or pain on CRF receptor 2 expression.

We also saw an increase in *crfr2* expression in the BLA following fear conditioning. Given the known role for this region in fear conditioning, this is not surprising. This is consistent with the critical role of the BLA in fear conditioning and fear expression. Previous work has also shown changes in CRF receptor signalizing and function during fear conditioning [67,72–75], although studies on CRFR2 are less consistent [76], and this work should be interpreted as broadly consistent with those findings.

The current data also demonstrate differences in CRF receptor expression in both the amygdala and hypothalamus as a function of hemisphere. While some of this was attributable to differences in ROI creation, there were several meaningful differences that this does not account for. For example, there were increases in CRF receptor 2 expression only in the right hemisphere of the BLA and CeA following early life pain and fear conditioning. This type of lateralization is likely reflective of our growing understanding that affective function is lateralized. In both pain and emotion processing, the left and right amygdala appear to have different functions with the right side being especially important for pain and fear responding [77–83]. In the PVN, the DAPI expression were higher on the left, but we found more *crhr1* expression (in terms of percent pixels) on the right. We also saw an effect of pain of *crhr2* expression, only on the left. While hemispheric specialization is less well understood in the hypothalamus, there is emerging evidence of lateralization there as well [84].

We also observed some sex differences in CRF receptor expression. This was most notable with greater *crhr2* in the male VMH, which is consistent with the known androgen-mediation of *crhr2* expression in this region [85]. We also observed female-specific changes in *crfr2* expression especially in the BLA, and less so in the CeA, following fear conditioning. The small effects of pain on the CeA were limited to males – consistent with our prior work [46,47]. Indeed, there are robust sex differences published in CRF signaling [86,87], stress [88–90], pain responding [91,92], HPA-axis activation [42], and the effects of early life stress and pain [91,93–98]. These effects have been associated with sex differences in CRF, ACTH and CORT expression. Thus, we anticipated sex differences in CRF receptor expression also.

There are some important limitations to this study. While we observed fear conditioning-induced differences in CRF receptor expression in the amygdala, suggesting that our timepoint (1 hour after onset of conditioning) was sensitive to this manipulation, many other studies have suggested that peak stress-induced expression is 1.5-4 hours after the stressor [99–101], although others have found increases earlier [102]. Thus, it’s possible that this study underestimates the effects of the fear conditioning manipulation on receptor expression. Of course, this limitation does not apply to any lasting changes that could have been induced by the neonatal manipulation, which was conducted weeks earlier. Second, RNAscope is designed to produce discrete puncta, with each dot representing a single transcript. Our imaging yielded a clear signal, but the degree of overlap and larger “blobs” did not leave us confident in our ability to assess individual puncta and complicated quantifying the degree of expression using individual dots as measures. This is common in studies using similar targets [103–105]. We dealt with this in two ways. First, we simply scored cells as having expression above threshold, or not. This does not depend on the resolution of discrete puncta. Second, we examined the percentage of pixels that expressed signal. This allows us to assess the degree to which clustering of multiple puncta occurred and likely reflects the degree of expression within cells already expressing the transcript. Our use of consistent imaging and thresholding parameters was designed to offset any potential confounds in this interpretation.

Overall, when combined with the literature, these data are consistent with the hypothesis that early life pain and juvenile stress alter CRF signaling in the amygdala and hypothalamus. While there are lasting effects of our NICU-model on CRF receptor expression, these appeared limited to CRF receptor 2 and predominately in the PVN. Our prior work demonstrating changes in expression of the CRF peptide itself, including the number of cells expressing CRF and amount of expression, is likely more important than long term changes in receptor distribution and expression, at least within in the amygdala [46,47]. The current data also demonstrate that CRF receptor expression is affected by juvenile fear conditioning in the amygdala, especially in females. Thus, we believe that early life pain and stress during a critical developmental period in the perinatal period alters development of CRF signaling creating an underlying and enduring vulnerability which alters later life responses to stress. It is likely the combination of these effects (early life trauma creating “organizational” changes and later life trauma “activating” those altered circuits) that likely creates the unique outcomes from both “hits” such as the tactile hypersensitivity we have observed in our previous work.

## Conclusions

Neonatal pain fails to alter of expression of CRF receptor 1. Our model did alter the expression of CRF receptor 2 mRNA in the hypothalamus. Fear conditioning primarily altered CRF receptor 2 in the amygdala of females. This has implications on the mechanisms by which early life trauma leads to a lasting vulnerability to a later life stressor or trauma.

